# Natural selection on synonymous genetic variation in the major histocompatibility complex

**DOI:** 10.64898/2026.02.23.707394

**Authors:** Jacob Roved

## Abstract

Protein coding DNA sequences harbor synonymous nucleotide variation that does not change amino acid sequences but influences phenotypes via multiple effects on the pathway from gene to protein. Synonymous variation has recently been shown to coevolve between viruses and their natural hosts, but its potential role in host immune defenses has not been explored. Here, I present evidence of natural selection on synonymous variation in the major histocompatibility complex (MHC), a highly polymorphic multigene locus that plays a crucial role in pathogen recognition by the adaptive immune system of vertebrate species.

Using data from a wild population of Great Reed Warblers, I show that codon usage in exon 3 of MHC class I (MHC-I) genes is under strong purifying selection in 57 out of 87 codon sites. Scanning the Great Reed Warbler genome for tRNA genes revealed that, for most amino acids, bias towards preferred codons was associated with abundances of tRNA isotypes, indicating that the purifying selection is likely driven by selection for increased translational efficiency. However, spikes of synonymous variation appeared in 30 of the 87 sites in the MHC-I exon 3, and in those sites, codon usage bias and correlations with tRNA abundances were reduced. About half of these sites fall in CpG contexts, where cytosine deamination elevates the mutation rate; the other half do not, leaving the driver of much of the variation unexplained. The distribution of the spikes of synonymous variation showed no consistent association with structural domains of the MHC-I protein, nor with sites under positive selection for amino acid change, which are considered important for antigen binding properties. Intriguingly, the amount of synonymous variation in genotypes showed a positive correlation with Darwinian fitness, indicating that important evolutionary forces are at play that neutralize purifying selection in the 30 sites.

From an ultimate perspective, the release of purifying selection among certain sites in MHC genes may indicate an arms race with pathogens, and I propose that the spikes of synonymous variation reveal a footprint of natural selection to escape inhibitory molecular interactions targeting MHC mRNA. I advance this RNA-level arms race as a novel axis of host–pathogen coevolution and a hypothesis to be tested directly. The fitness association is specific to synonymous variation, independent of amino-acid differences and the number of MHC alleles in genotypes, implicating selection on the synonymous variation itself rather than linked protein-level diversity. These results expose a previously unrecognized layer of selection in a key immune gene family and highlight the potential functional importance of synonymous variation in host–pathogen coevolution with direct implications for understanding susceptibility to infectious diseases.

**Significance Statement:** Roughly a quarter of DNA mutations carry no amino acid change, yet a growing body of evidence shows such synonymous variation can still shape how genes work. This study shows that genes of the major histocompatibility complex, central to how vertebrates detect infection, carry a hidden layer of natural selection acting on this synonymous variation, and that it is linked to individual fitness in a wild bird population. The findings suggest hosts and pathogens may be locked in an evolutionary arms race playing out beneath the level of protein sequence, with implications for how susceptibility to infectious disease is understood and studied.

## Introduction

A much-overlooked type of natural selection is that which concerns synonymous nucleotide variation that does not change amino acid sequences. Approximately one quarter to one third of single nucleotide mutations are expected to be synonymous and were previously presumed to have little or no effects on phenotypes (*1–3*). However, it is now generally acknowledged that synonymous nucleotide variation can cause changes in protein expression, conformation, and function, and may be subject to both purifying and positive selection (*3–7*).

Codon usage bias (CUB) is the common term describing the unequal use of synonymous codons in coding genomic regions (*3*, *5*, *7*). CUB is prevalent in highly expressed genes especially in species with large population sizes and is thought to be driven by optimization of codon usage to the availability of tRNA isotypes. Preferred codons are favored by natural selection because they speed up mRNA translation by ribosomes and reduce the risk of mistranslation, where incorporation of a different amino acid could introduce potentially detrimental changes to the protein (*3*, *8*, *9*). However, optimization for fast expression is balanced by other factors, notably co-translational protein folding, especially in higher organisms. Codon usage can slow the progression of the ribosome, through rare-tRNA codons or stable local mRNA structures, giving a nascent protein domain time to fold without interference from downstream elements; accordingly, slow-folding β-sheets and interdomain bridges harbor more rare codons than fast-folding α-helices (*3*, *5*, *6*, *10*, *11*). Furthermore, codon usage at specific sites may be conserved by molecular interactions, e.g., of DNA with transcription factors or of mRNA with microRNAs, splicing enhancers, deaminases, ribosomes, and other molecules that regulate transcription and translation (*3*, *6*, *7*, *12*, *13*).

Following *the nearly neutral theory*, it has been predicted that synonymous SNPs are unlikely to be affected by natural selection in species that have small population sizes, such as many vertebrate species (*13*, *14*). That prediction rests on the assumption that synonymous variation is mostly under weak selection, which appears to be the general trend observed across genomes, e.g., in a comparison of 5,639 orthologous genes between human, chimpanzee, macaque, mouse, and rat (*4*). However, some loci evolve under abnormal regimes of natural selection. Numerous disease associations involving synonymous SNPs have been detected and a recent study estimated that 25.9% of synonymous SNPs are under weak and 3.6% under strong negative selection in humans (*6*, *15*, *16*). A particularly interesting candidate locus for effects of synonymous nucleotide variation in vertebrate species is the major histocompatibility complex (MHC). The genes of the MHC encode molecules that present peptide antigens to T-cells, which is a decisive step in the induction of adaptive immune responses (*17*, *18*). MHC genes are engaged in a coevolutionary arms race with pathogens that has generated both a great diversity in these genes and preservation of polymorphisms over evolutionary time (*17*, *19–23*).

Among MHC studies, a broad consensus has prevailed to regard synonymous nucleotide variation as silent and to regard variation in protein sequences and gene expression as the only factors determining phenotypic variation. However, biological effects of synonymous nucleotide variation in the MHC have never been thoroughly investigated, leaving a major gap in our knowledge about a substantial body of genetic variation in these genes that serve a central function in vertebrate immune systems. The single prior characterization of synonymous variation in an MHC gene reported conspicuous non-random “spikes” of synonymous variation in the human HLA-A locus, but attributed them to a neutral mutational process, cytosine deamination, without testing whether they reflect selection (*24*). Whether such site-specific synonymous variation in the MHC represents neutral mutational artefacts or signatures of selection has therefore remained unresolved. Protein sequence variation in the MHC has been shown to affect the binding of antigen peptides and to be under positive selection, and it is evident that this variation is of evolutionary importance (*22*, *23*, *25*, *26*). However, the assumption that synonymous variation in the MHC serves no biological function involves a severe risk of accumulating wrong or biased insights.

A recent study observed that CUB in viruses tended to be more similar to that of symptomatic hosts than that of natural hosts (i.e., hosts that the virus coevolved with). The authors showed that expression of virus genes with CUB similar to the host severely impeded translation of host genes by depletion of tRNAs used by virus genes, supporting a general deleterious effect of CUB similarity (*27*). Furthermore, infectious pathogens have evolved specific strategies to manipulate host immune systems to evade detection, including molecular interactions that target host DNA and mRNA to disrupt gene expression and protein synthesis in key immune system pathways (*28–33*). This previously unrecognized complexity in host-pathogen coevolution invites the question of whether antagonistic coevolution of codon usage between viruses and hosts may affect MHC genes, the expression of which is crucial to adaptive immune responses of vertebrate hosts. The fitness advantage to pathogens that evade host immunity are obvious and exemplified e.g. by convergent evolution among several viruses of mechanisms that target the mRNA of the MHC class I polypeptide-related sequence B (MICB) gene (*34–36*). Such interventions from pathogens are likely to elicit evolutionary responses in the host, but host immunity genes are evolutionarily constrained by the function of the encoded proteins. Synonymous mutations generate a substantial body of genetic variation that can engage in evolutionary responses to natural selection from pathogens independent from this constraint (*6*, *10*, *27*). If hosts become more often or more severely infected by inhibition of MHC expression by pathogens, then synonymous nucleotide variation may be subject to an arms race, where the aim of the host would be to escape regulatory mechanisms of pathogens to ensure gene expression.

Here I characterized for the first time synonymous nucleotide variation in the MHC of a wild vertebrate species and investigated whether it is subject to natural selection. I used data from a 20-year study of a wild breeding population of Great Reed Warblers *Acrocephalus arundinaceus*, a socially polygynous, migratory songbird. I employed 559 genotypes of the highly polymorphic MHC class I (MHC-I) exon 3, which encodes a major section of the antigen binding groove in MHC-I molecules (*37*), along with the Great Reed Warbler genome assembly and field observations of individual life histories and lifetime reproductive success (*25*, *38–40*). This detailed ecological data set provides a unique opportunity to combine analyses of synonymous nucleotide variation in the MHC with investigations of direct effects on individual Darwinian fitness.

Specifically, I tested whether synonymous codon usage among 390 Great Reed Warbler MHC-I exon 3 sequences showed evidence of natural selection and analyzed how synonymous nucleotide variation differs between sites along the exon. I compared site-specific synonymous variation with sites predicted to be under positive selection for amino acid changes and with predicted structural domains of the folded MHC-I protein and tested whether it could be explained by cytosine deamination by examining the sequence context of variable sites. Additionally, I employed a novel approach that compared site specific variation in the 390 empirical MHC-I exon 3 sequences with 1,000 synonymous data sets simulated under the assumption of no selection on codon usage. Furthermore, I quantified CUB and analyzed whether codon usage in the MHC-I is correlated with frequencies of cognate tRNA isotypes in the Great Reed Warbler genome, a proxy for tRNA availability (*10*). Finally, I calculated the mean number of synonymous nucleotide changes among the MHC-I sequences in each individual and analyzed whether synonymous nucleotide variation in the Great Reed Warbler MHC-I was associated with life span or Darwinian fitness.

## Results

### Selection analyses

I employed mutation-selection models in codeml (*41*, *42*) to test the null hypothesis that variation in codon usage in exon 3 of MHC-I genes (Table 1) is shaped by mutation bias alone and not by selection on synonymous nucleotide variation, cf. (*4*). I have previously demonstrated that the Great Reed Warbler MHC-I exon 3 sequences show evidence of positive selection for amino acid changes (*25*), and I therefore nested mutation-selection and null-models within the site model M8, which allows a category of sites evolving under positive selection (Ω > 1, where Ω is the dN/dS ratio) (*4*, *43*). The mutation-selection model (FMutSel), reflecting the alternative hypothesis that codon usage is affected by selection, fit the data significantly better than the null-model (FMutSel0), which assumes that codon usage is only affected by mutation bias (likelihood ratio test: p = 2.43 × 10^-15^; Table 2). Because FMutSel estimates codon fitnesses that are fixed and shared across all sites, this improved fit identifies directional selection consistently favoring particular codons over their synonyms (i.e., CUB), rather than selection that maintains synonymous diversity. On balance the favored codons are G/C-ending: within-block codon usage is shifted toward G/C-ending codons relative to the mutation-only expectation in 16 of 18 synonymous blocks (paired t-test, t = 5.48, d.f. = 17, p = 4.1 × 10⁻⁵; Table S1), whereas the mutational process inferred by the same model is strongly A/T-biased (Table S2; per-codon values in Table S3). The inferred selection on codon usage therefore opposes the mutational bias.

**Table 1.**
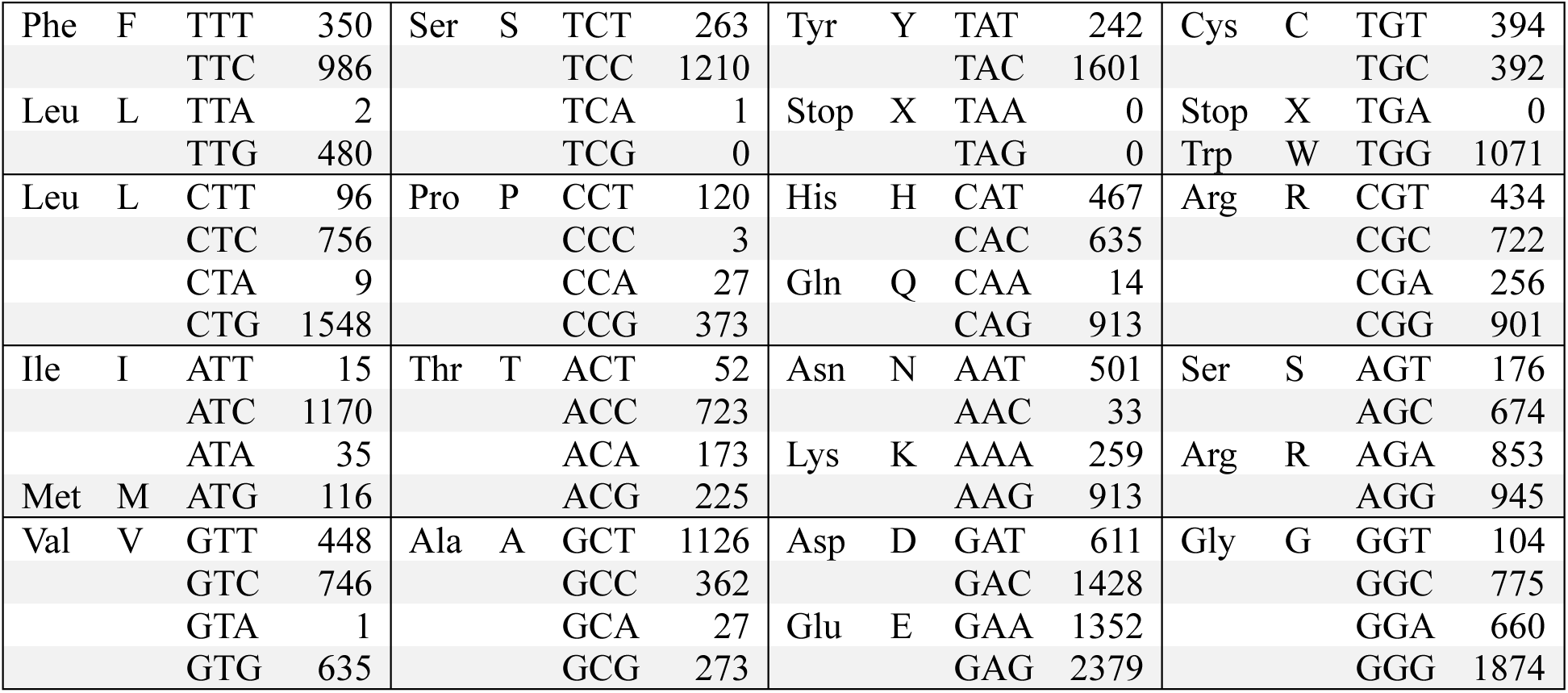
Codon usage counts among the 390 Great Reed Warbler MHC-I exon 3 sequences.

**Table 2.** Summary of the M8 mutation-selection vs. null models in codeml. ln(L) specifies the log likelihood of the models, Ω > 1 specifies the proportion of sites that were estimated to be under positive selection for amino acid change, and Ω est indicates the value of Ω estimated for that category of sites. The p-value was calculated by likelihood ratio test of the nested FMutSel vs. FMutSel0 models.

| **Codon model** | **Hypothesis** | **ln(L)** | **Ω > 1** | **Ω est** | **p(H<sub>0</sub>)** |
|---|---|---|---|---|---|
| FMutSel0 | H<sub>0</sub>: No selection on synonymous variation | -7521.6 | 21.2% | 4.22 | 2.43 × 10<sup>-15</sup> |
| FMutSel | H<sub>1</sub>: Selection on synonymous variation | -7443.6 | 15.5% | 3.23 |  |

Notably, the null-model overestimated both the proportion of sites with Ω > 1 and the Ω estimate for that category, compared to the mutation-selection model (Table 2). This shows that ignoring selection on synonymous codon usage inflates estimates of positive selection on amino acid variation in codeml. Repeating the analysis using the site model M2a confirmed the results from the M8 model (Table S4). Sites with Ω > 1 inferred by Bayes Empirical Bayes analysis in the M8 mutation-selection model are specified in Table S5 (44).

### Patterns of synonymous variation

To further investigate the patterns of synonymous nucleotide variation, I quantified the synonymous changes per base and per codon site in pairwise comparisons of the 390 MHC-I exon 3 sequences. The frequency of synonymous codon variation differed greatly along the sequences, with most sites showing little or no synonymous variation, but with prominent spikes appearing in roughly one third of the sites along the sequence (sites 1, 3, 5, 11, 12, 14, 17, 18, 23, 24, 31, 35, 38, 40, 41, 53, 54, 57, 61, 63, 66, 68, 69, 71, 72, 74, 77, 78, 80, and 85; N=30; Fig. 1a). The synonymous codon variation was mostly associated with variation in the third nucleotide position, except in site 61, where more than half of the observed synonymous variation involved a change in the first nucleotide position (Table S6). That was also the case for site 33, but that site harbored almost no synonymous variation (freq. ∼0.005). There was no consistent association between frequencies of synonymous codon variation and the location of encoded amino acids in bridge domains, β-sheets, or α-helices of the protein structure predicted with AlphaFold 3 (*45*) (Fig. 1a; Fig. S1; Table S7). Sites that were inferred to evolve under positive selection for amino acid change (i.e., sites with Ω > 1 marked by asterisks in Fig. 1a) showed no obvious trend towards harboring high or low synonymous variation.

**Fig. 1.**
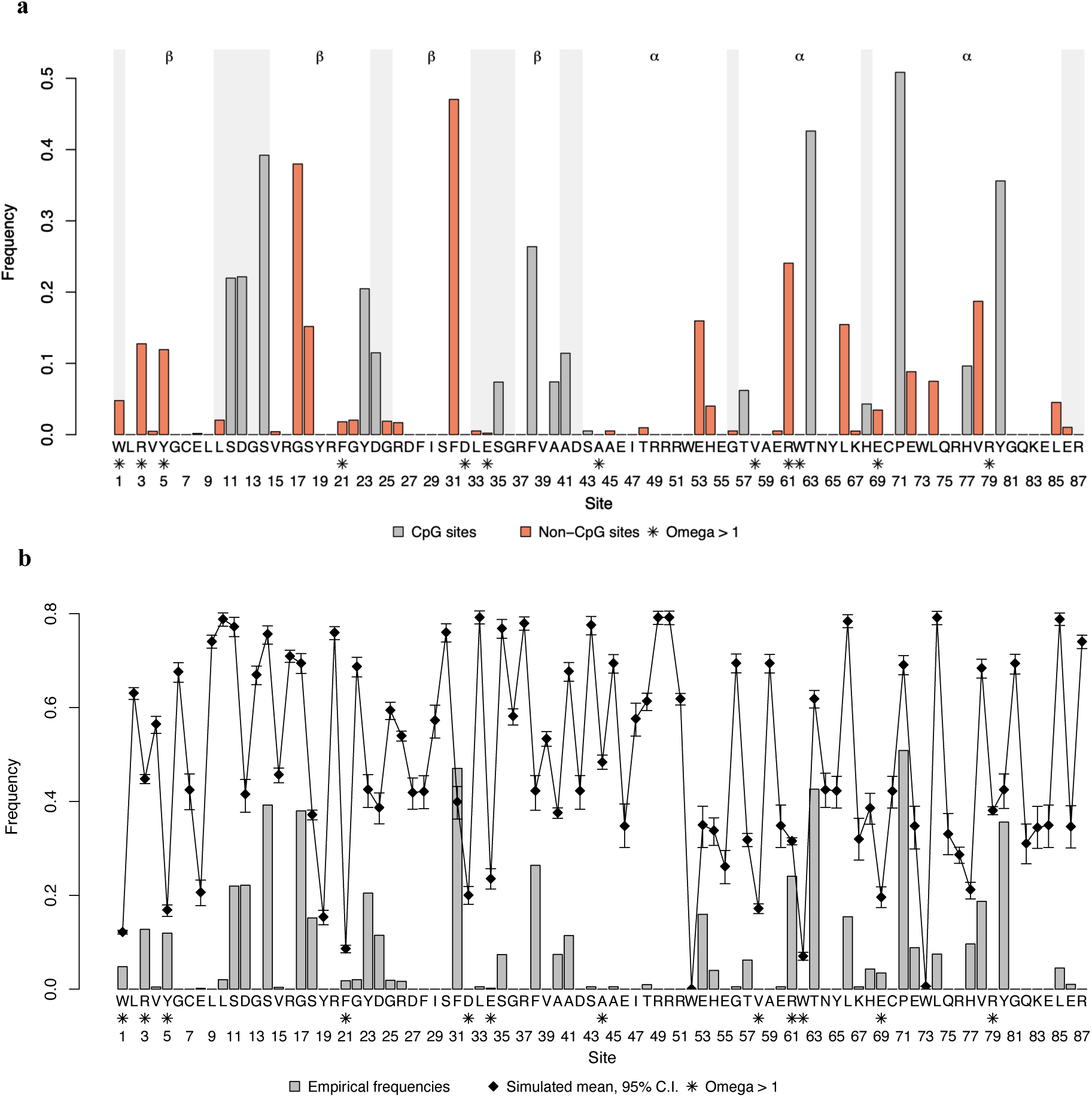
Frequency of synonymous changes for each amino acid site among the 390 Great Reed Warbler MHC-I exon 3 sequences. **(a)** Gray bars indicate codons whose third position lies in a CpG dinucleotide; orange indicates non-CpG codons. Shaded areas mark bridges between α-helix and β-sheet domains in the predicted protein structure. **(b)** Bars indicate the frequency of synonymous changes for each site in the empirical data set (same as the bars in a). Black diamonds indicate the mean frequency of synonymous changes for each site among 1,000 data sets simulated under the assumption of no selection, with error bars indicating 95% C.I. The consensus amino acid sequence indicates the most frequent amino acid for each site. Asterisks mark the sites predicted to evolve under positive selection for amino acid changes by Bayes Empirical Bayes analysis in the M8 FMutSel model.

To understand how selection has shaped synonymous nucleotide variation along the MHC-I exon 3 sequences, I simulated 1,000 data sets of 390 nucleotide sequences synonymous to the empirical Great Reed Warbler MHC-I exon 3 sequences (i.e., with identical amino acid sequences), assuming the null hypothesis that variation in codon usage is due to mutation bias alone and not affected by selection acting at synonymous codons. The frequencies of synonymous variation among the sequences in the simulated data sets highlight the non-random distribution of synonymous variation among the empirical MHC-I sequences: 86 out of the 87 sites harbored significantly less synonymous differences than predicted from the sequences simulated under the assumption of no selection, providing a further indication of purifying selection on codon usage in the Great Reed Warbler MHC-I (Fig. 1b; Table S8). However, the spikes of synonymous variation along the sequences indicate that the purifying selection is relaxed in about one third of the sites. Site 31 showed increased synonymous variation compared to the simulated data sets (p < 0.001; Fig. 1b; Table S8), but it contains a near-balanced phenylalanine (Phe) TTT/TTC polymorphism with a hypermutable cytosine, so this excess must be interpreted with caution (Table S9).

The simulations above model mutation bias at single nucleotides and do not capture context-dependent mutation. Matsushita & Kano-Sueoka attributed the analogous non-random synonymous variation in human HLA-A to cytosine deamination, a neutral mutation-pressure process (*24*). I therefore tested whether deamination explains the spikes in the Great Reed Warbler MHC-I, examining the CpG dinucleotide context. Spikes and conserved sites did not differ in the proportion of third positions occupied by a cytosine or guanine, the substrate for deamination (22/30 vs. 43/57; Fisher’s exact test, p = 1; Table S10). The spikes were, however, strongly enriched for cytosines in a CpG dinucleotide context (the most deamination-prone context, where methylated cytosines deaminate at elevated rates): half of the spikes (15/30) fell in a CpG context, versus a single site under strong purifying selection (1/57; odds ratio ≈ 53, Fisher’s exact test, p = 7.6 × 10⁻⁸; Fig. 1a; Table S10). Cytosine deamination at CpG sites is therefore a major driver of where the spikes occur. It cannot, however, account for all of them: the other half of the spikes (15/30) occur outside CpG contexts, and their elevated synonymous variation is not explained by CpG hypermutability.

### Codon usage bias

For most degenerate amino acids, CUB was greater in the 390 Great Reed Warbler MHC-I exon 3 sequences than in the 1,000 data sets simulated under the assumption of no selection on synonymous codon usage (Fig. 2; Table S11). The exceptions are histidine (His), glutamic acid (Glu), cysteine (Cys), and arginine (Arg), where the observed CUBs were smaller than expected, and lysine (Lys) and aspartic acid (Asp), where the observed CUBs were similar to those in the simulated data sets. CUB was on average greater among the sites where synonymous variation is under purifying selection compared to the sites where selection is relaxed (paired t-test, t = 2.22, d.f. = 16, p = 0.021). However, for Leu, Val, Asp, Arg, and glycine (Gly) it was smaller (Fig. 3; Table S12).

**Fig. 2.**
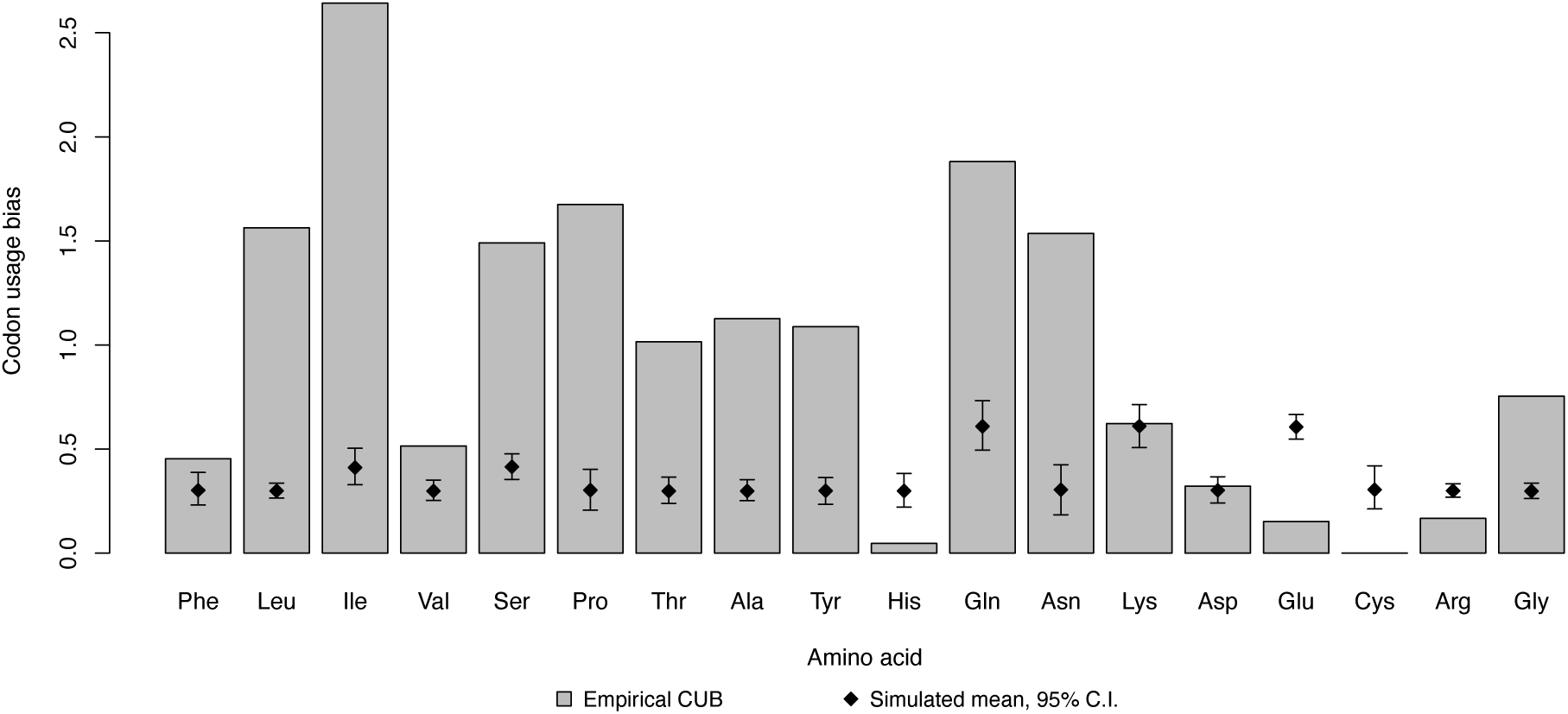
Codon usage bias (CUB) measured as the variance in relative synonymous codon usage for degenerate amino acids. Gray bars show the CUBs observed among the 390 empirical MHC-I exon 3 sequences. Black diamonds show the mean CUBs observed among 1,000 data sets simulated under the assumption of no selection on synonymous codon usage, with error bars indicating 95% C.I.

**Fig. 3.**
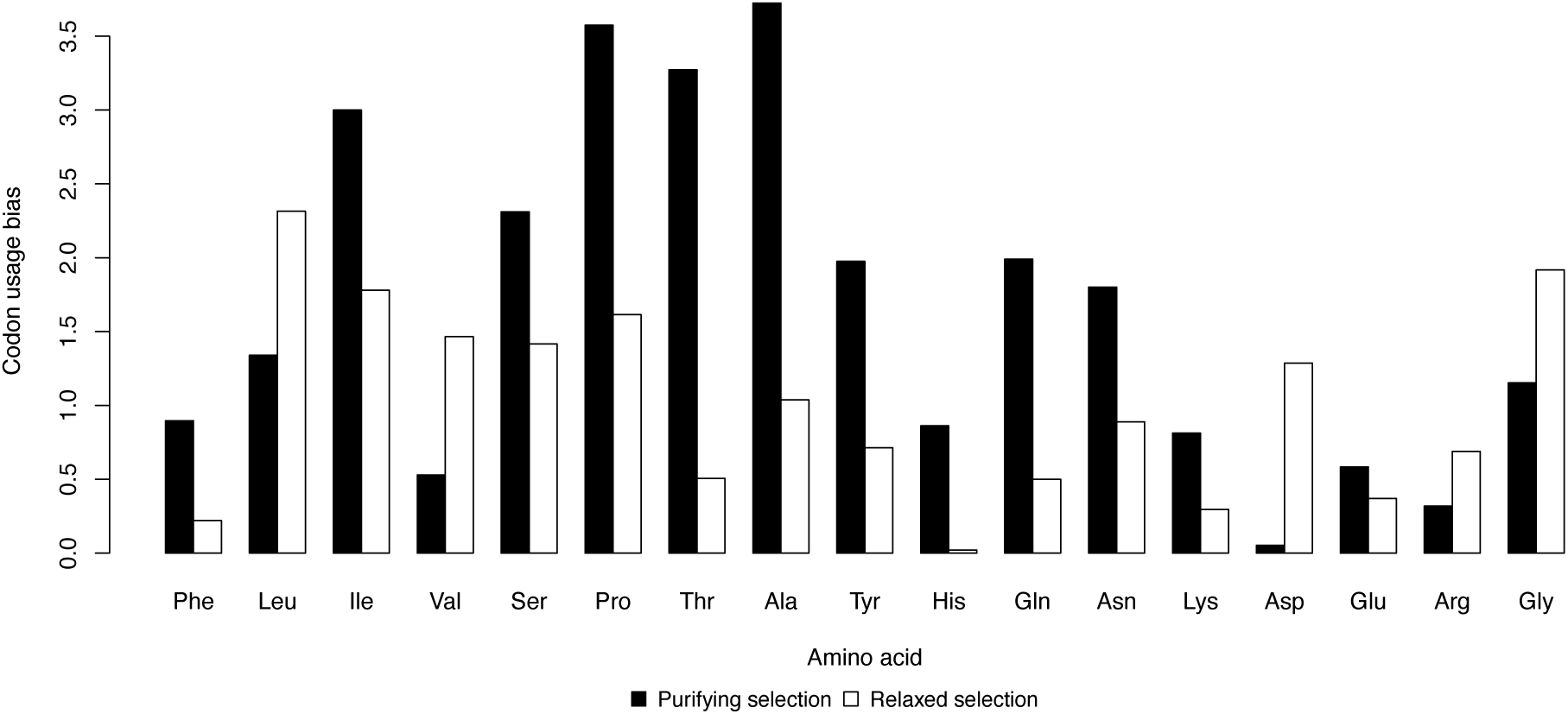
Codon usage bias (CUB) measured as the variance in relative synonymous codon usage for degenerate amino acids among the 390 empirical MHC-I exon 3 sequences. Black bars show the CUBs measured in the 57 sites that evolve under purifying selection for codon usage. White bars show the CUBs measured in the 30 sites where purifying selection for codon usage was relaxed.

### Codon usage and abundance of tRNA isotypes

To estimate tRNA isotype availability, I employed tRNAscan-SE (*46*) to scan the Great Reed Warbler genome for tRNA genes. The analysis detected 325 functional genes for cytosolic standard amino acid tRNAs. The counts of tRNA isotypes among the 325 genes are shown in Table 3. Fourteen codons encoding 13 amino acids were not matched by a tRNA isotype in the genome. Twelve amino acids were matched by more than one tRNA isotype in the Great Reed Warbler genome. For seven of those amino acids (Leu, Val, serine (Ser), alanine (Ala), glutamine (Gln), Lys, and Glu), relative synonymous codon usage (RSCU) in the 390 MHC-I exon 3 sequences was positively correlated with normalized relative frequencies of synonymous tRNA isotypes (Fig. 4; Table S13; S14). For Leu, Val, Ser, and Ala the correlation coefficients were significantly more positive than expected from comparisons with sequences simulated under the assumption of no selection, while for Gln, Lys, and Glu the correlation coefficients were 1 with both the empirical and simulated data sets.

**Fig. 4.**
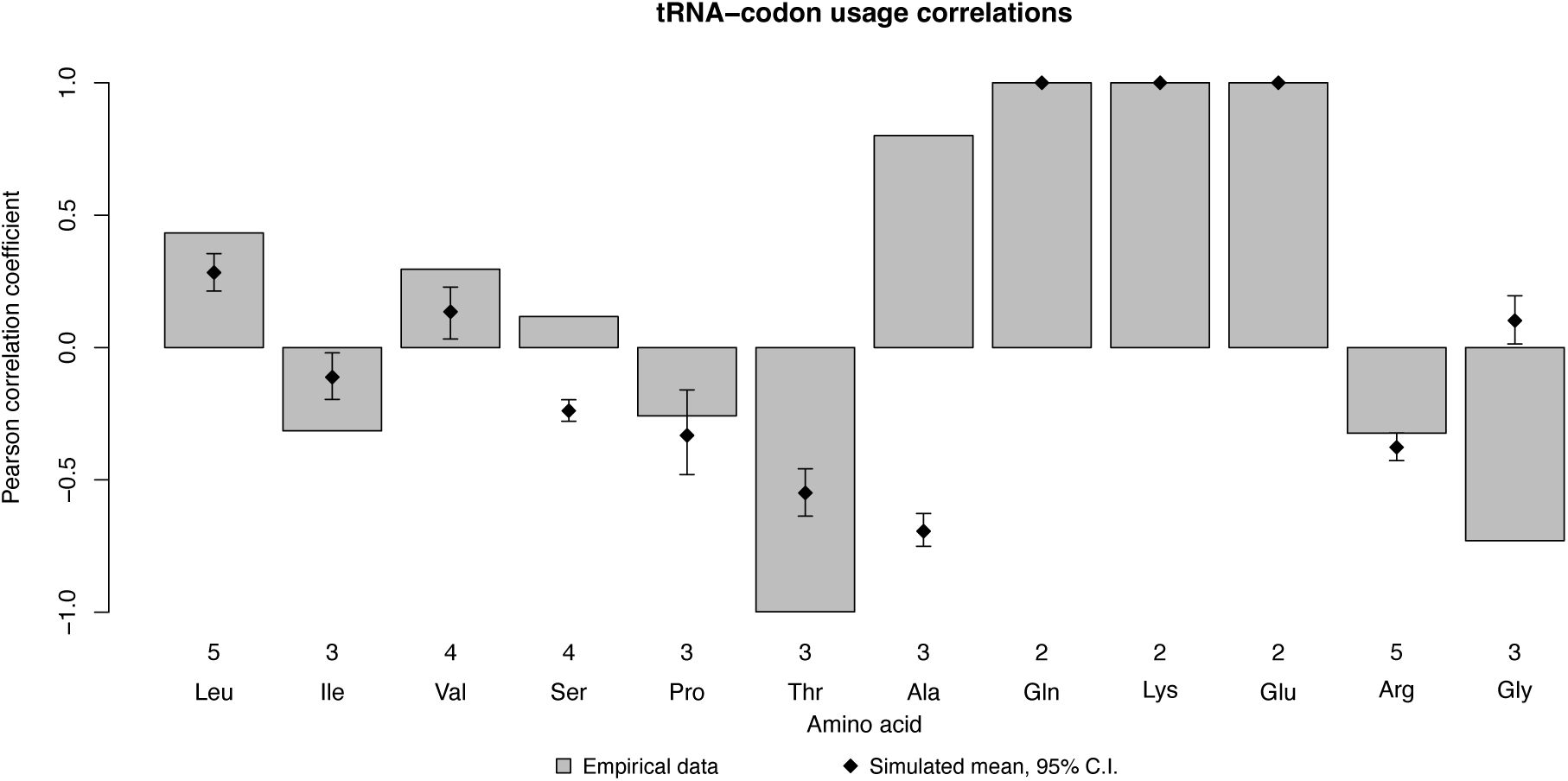
The relationship between normalized relative frequencies of synonymous tRNA isotypes in the Great Reed Warbler genome and relative synonymous codon usage (RSCU). Gray bars show Pearson correlation coefficients for RSCU in the 390 empirical MHC-I exon 3 sequences. Black diamonds show the mean Pearson correlation coefficients for RSCU among 1,000 data sets simulated under the assumption of no selection on synonymous codon usage, with error bars indicating 95% C.I. The numbers under the bars specify the number of codons matched by tRNA isotypes for each amino acid. Only amino acids with more than one tRNA isotype in the genome were included in the analyses.

**Table 3.** Isotype counts among the 325 tRNA genes in the Great Reed Warbler genome. Anticodons are specified in italics.

|  |  |  |  |  |  |  |  |  |  |  |  |  |  |  |  |  |  |  |  |
|---|---|---|---|---|---|---|---|---|---|---|---|---|---|---|---|---|---|---|---|
| Phe | F | TTT | *AAA* | 0 | Ser | S | TCT | *AGA* | 12 | Tyr | Y | TAT | *ATA* | 0 | Cys | C | TGT | *ACA* | 0 |
|  |  | TTC | *GAA* | 7 |  |  | TCC | *GGA* | 0 |  |  | TAC | *GTA* | 9 |  |  | TGC | *GCA* | 28 |
| Leu | L | TTA | *TAA* | 1 |  |  | TCA | *TGA* | 3 | Stop | X | TAA | *TTA* | 0 | Stop | X | TGA | *TCA* | 0 |
|  |  | TTG | *CAA* | 2 |  |  | TCG | *CGA* | 6 |  |  | TAG | *CTA* | 0 | Trp | W | TGG | *CCA* | 5 |
| Leu | L | CTT | *AAG* | 6 | Pro | P | CCT | *AGG* | 7 | His | H | CAT | *ATG* | 0 | Arg | R | CGT | *ACG* | 6 |
|  |  | CTC | *GAG* | 0 |  |  | CCC | *GGG* | 0 |  |  | CAC | *GTG* | 7 |  |  | CGC | *GCG* | 0 |
|  |  | CTA | *TAG* | 2 |  |  | CCA | *TGG* | 3 | Gln | Q | CAA | *TTG* | 3 |  |  | CGA | *TCG* | 2 |
|  |  | CTG | *CAG* | 5 |  |  | CCG | *CGG* | 3 |  |  | CAG | *CTG* | 12 |  |  | CGG | *CCG* | 1 |
| Ile | I | ATT | *AAT* | 46 | Thr | T | ACT | *AGT* | 6 | Asn | N | AAT | *ATT* | 0 | Ser | S | AGT | *ACT* | 0 |
|  |  | ATC | *GAT* | 12 |  |  | ACC | *GGT* | 0 |  |  | AAC | *GTT* | 7 |  |  | AGC | *GCT* | 5 |
|  |  | ATA | *TAT* | 2 |  |  | ACA | *TGT* | 3 | Lys | K | AAA | *TTT* | 6 | Arg | R | AGA | *TCT* | 3 |
| Met | M | ATG | *CAT* | 15 |  |  | ACG | *CGT* | 2 |  |  | AAG | *CTT* | 10 |  |  | AGG | *CCT* | 3 |
| Val | V | GTT | *AAC* | 4 | Ala | A | GCT | *AGC* | 15 | Asp | D | GAT | *ATC* | 0 | Gly | G | GGT | *ACC* | 0 |
|  |  | GTC | *GAC* | 1 |  |  | GCC | *GGC* | 0 |  |  | GAC | *GTC* | 7 |  |  | GGC | *GCC* | 12 |
|  |  | GTA | *TAC* | 1 |  |  | GCA | *TGC* | 8 | Glu | E | GAA | *TTC* | 4 |  |  | GGA | *TCC* | 7 |
|  |  | GTG | *CAC* | 4 |  |  | GCG | *CGC* | 3 |  |  | GAG | *CTC* | 5 |  |  | GGG | *CCC* | 4 |

When comparing the 57 sites that evolve under purifying selection for synonymous codon usage with the 30 sites where purifying selection for codon usage was relaxed, a different image appeared. Overall, the correlation coefficients were significantly greater among the 57 sites that evolved under purifying selection than among the 30 sites where selection was relaxed (paired t-test, t = 2.05, d.f. = 11, p = 0.032, mean diff. = 0.63; Fig. 5; Table S15). The amino acids Val, threonine (Thr), Ala, Lys, and Glu showed strong positive correlations among the 57 sites that evolve under purifying selection for codon usage, but negative correlations among the 30 sites where purifying selection for codon usage was relaxed. A contrasting pattern was also seen for Ser, where the correlation was negative among the 57 sites and positive among the 30 sites. Leu, isoleucine (Ile), proline (Pro), Arg, and Gly showed correlations in similar directions between the 57 sites and the 30 sites, although with different strength. For Gln the correlation coefficients were 1 among both the 57 sites and the 30 sites. Together, these observations suggest that synonymous codon usage in the Great Reed Warbler MHC-I is affected by availability of cognate tRNAs during translation, and that the associations differ both between amino acids and between sites.

**Fig. 5.**
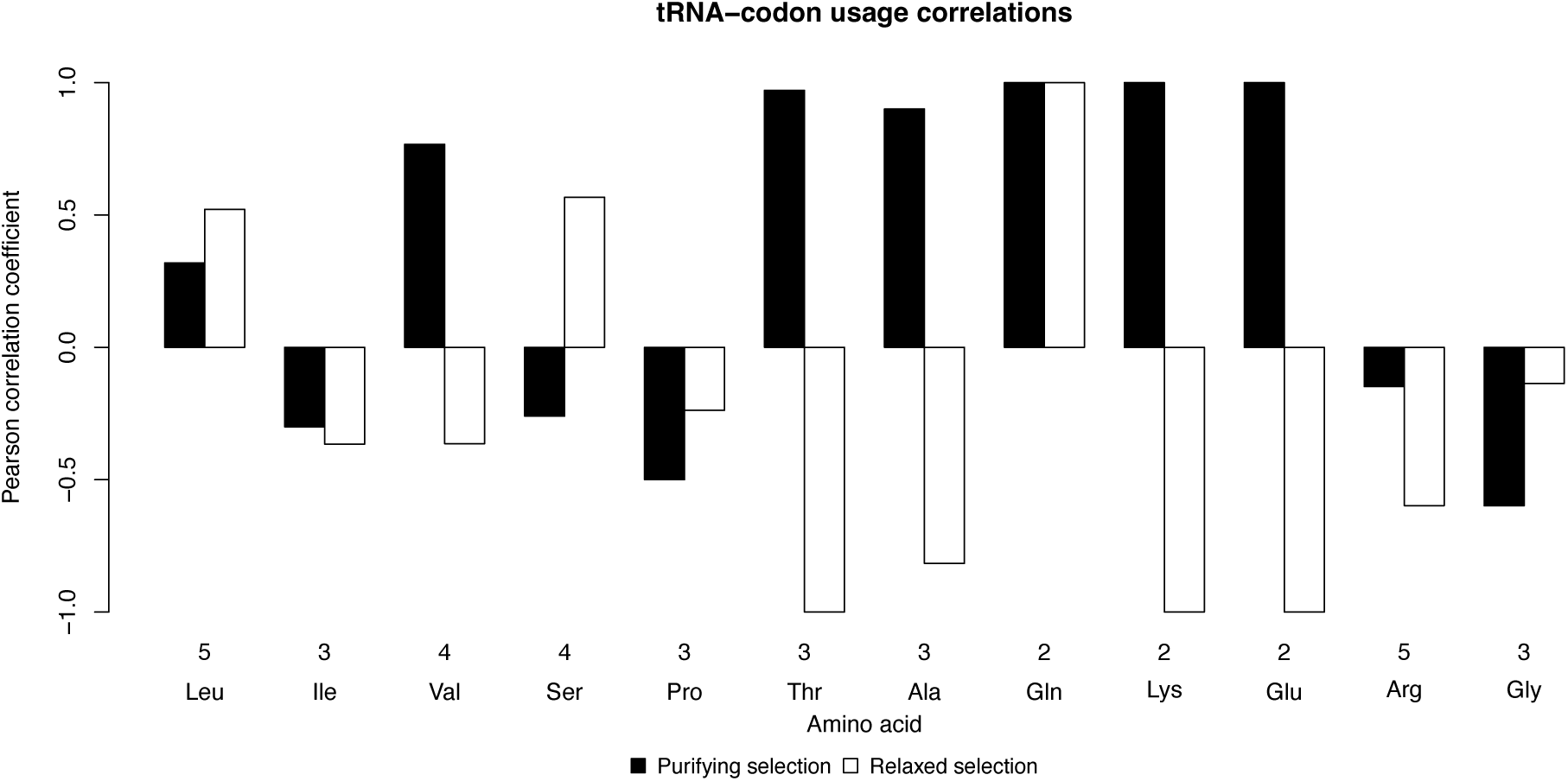
The relationship between normalized relative frequencies of synonymous tRNA isotypes in the Great Reed Warbler genome and relative synonymous codon usage (RSCU). Black bars show Pearson correlation coefficients for RSCU calculated among the 57 sites of the empirical 390 MHC-I exon 3 sequences that evolve under purifying selection for codon usage. White bars show Pearson correlation coefficients for RSCU calculated among the 30 sites where selection for codon usage was relaxed. The numbers under the bars specify the number of codons matched by tRNA isotypes for each amino acid. Only amino acids with more than one tRNA isotype in the genome were included in the analyses.

### Fitness effects

To investigate whether synonymous nucleotide variation in the MHC-I has a measurable effect on the survival or fitness of individuals, I employed life history data from the long-term study of Great Reed Warblers at Lake Kvismare in Sweden (*38*). For statistical analyses of fitness effects, I calculated the mean number of synonymous nucleotide changes in pairwise comparisons among the exon 3 sequences across all targeted MHC-I loci in each individual. The mean number of synonymous nucleotide changes in each individual ranged from 4.90 to 8.48 and followed a normal distribution with mean = 6.69 and s.d. = 0.64 (Fig. S2).

Among male Great Reed Warblers, the mean number of synonymous nucleotide changes in each individual was positively associated with life span (glm, b = 0.18, t = 2.21, p = 0.030, d.f. = 76; Fig. 6a; Table S16) and lifetime number of fledged offspring (glm, b = 0.36, t = 2.33, p = 0.022, d.f. = 76; Fig. 6b; Table S17). However, no association was observed in females (*life span*: glm, b = -0.058, t = -0.61, p = 0.54, d.f. = 95; Fig. 6a; Table S16; *lifetime number of fledged offspring*: glm, b = -0.015, t = -0.13, p = 0.90, d.f. = 95; Fig. 6b; Table S17). As common in species with discrete breeding events, lifetime fitness was strongly associated with life span in the Great Reed Warbler. I therefore also analyzed the effects of the mean number of synonymous nucleotide changes on lifetime number of fledged offspring in a model that adjusted for individual life span. In this model, the mean number of synonymous nucleotide changes in each individual showed a trend towards a positive effect in both males and females (glm, b = 0.10, t = 1.77, p = 0.079, d.f. = 172; Fig. S3; Table S18).

**Fig. 6.**
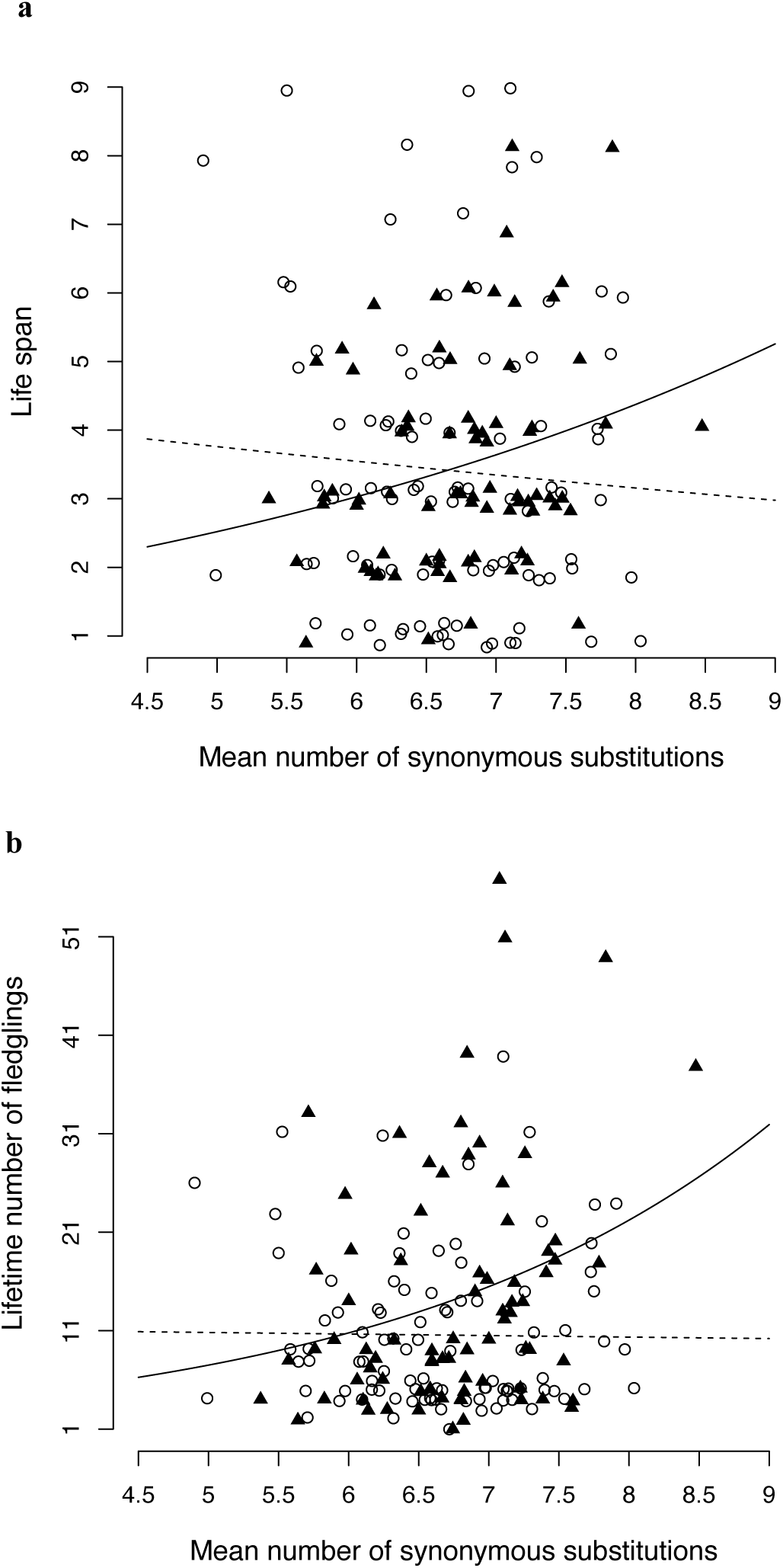
The relationship between synonymous nucleotide variation and **(a)** life span and **(b)** lifetime number of fledged offspring in adult Great Reed Warblers. The lines show predictions from generalized linear models that included sex and the interaction ‘mean number of synonymous nucleotide changes × *sex’. Open circles and dashed lines indicate females. Black triangles and solid lines indicate males. Jitter was added to life span and lifetime number of fledglings to distinguish discrete data points*.

No effects were observed of the number of different MHC-I sequences or mean amino acid p-distance between the MHC-I sequences in each individual. These variables were included as covariates in the full models but removed by the model simplification step in each analysis. Results from models of the effects of number of different MHC-I sequences and mean amino acid p-distance on life span, lifetime number of fledged offspring, and lifetime number of fledged offspring adjusted for life span are presented in Tables S19–S21.

## Discussion

### Selection on synonymous codon usage

This study shows for the first time that MHC genes are subject to natural selection on synonymous nucleotide variation and that this variation has direct measurable effects on Darwinian fitness of individuals. Intriguingly, 57 sites in the Great Reed Warbler MHC-I exon 3 showed evidence of strong purifying selection on codon usage, while selection was relaxed in the remaining 30 sites (Fig. 1a, b). Both CUB and the correlations between CUB and abundances of cognate tRNA isotypes in the genome were on average greater in the sites under strong purifying selection than in the sites where selection was relaxed, indicating that selection for translation speed and fidelity may generally restrict codon usage in the Great Reed Warbler MHC (Fig. 3; Fig. 5). The driving mechanisms behind such an effect may be optimization of codons to the availability of aminoacyl-tRNAs for translation and the advantage of recycling ribosomes (*3*, *7*, *9*, *47*).

Some other forces must be at play that allows selection to be relaxed in the remaining 30 sites. A possible explanation is that unpreferred codons may temporarily slow translation, allowing time for the nascent protein to fold without interference from downstream elements (*11*), but the distribution of the sites with higher frequencies of synonymous variation showed no consistent association with predicted domains of the folded MHC-I protein (Fig. 1a). However, the observed patterns warrant further investigations, as three of the four interdomain bridges following β-sheets in the first half of the exon showed elevated frequencies of synonymous variation. Temporary slowing of translation may not just be caused by use of rare tRNAs but may also be enforced by local stable mRNA structures that physically slow the progression of the ribosome. Synonymous nucleotide variation is known to affect mRNA structure and stability (*5*), but it is unclear how selection for local mRNA structures would affect its distribution in relation to the encoded protein domains. This is an interesting avenue for future investigations, but it is beyond the scope of the present study as proper modelling of the mRNA structure would require long range transcripts of complete MHC genes, which have proven extremely challenging to obtain on a large scale even with long-range sequencing technologies (*48–54*). There was also no apparent association between the distribution of synonymous nucleotide variation and sites that were inferred to evolve under positive selection for amino acid change (i.e., sites with Ω > 1). However, a recent study found that the Great Reed Warbler MHC-I binds antigen in a flat conformation due to a tweezer-like interaction between two arginines that form a conserved restriction point within the peptide binding groove (*55*). Interestingly, both sites forming this restriction point show elevated synonymous variation (sites 3 and 61; Fig. 1a).

The observed correlations with individual survival and fitness in the study population indicate that synonymous nucleotide variation in the MHC-I is of significant biological importance (Fig. 6). The fitness effects of synonymous variation were not accompanied by effects of number of different MHC-I sequences in each individual or amino acid distance between sequences, implicating synonymous variation specifically as the signal carrier. The effects observed on life span and lifetime number of fledglings represent a stronger influence on individual fitness than the previously reported effects of number of different MHC-I sequences on offspring recruitment success (*38*). The positive correlation with life span suggests that the phenotypic advantage of high synonymous nucleotide variation in males is largely associated with survival, however the trend towards a positive effect on lifetime number of fledged offspring in a model that adjusted for life span suggested that both males and females with high synonymous variation may on average rear more offspring per breeding year (Fig. S3). The synonymous nucleotide changes among the MHC-I sequences in each individual were quantified as an average distance across all targeted MHC-I loci, ruling out potential confounding effects of variation in other linked genes.

One concern is that the mutation–selection models capture an unmodelled mutational process, such as cytosine deamination at CpG sites, rather than selection. However, the model itself resolves this: FMutSel infers a strongly A/T-biased mutational spectrum consistent with deamination, yet the codon preferences it favors are G/C-ending and therefore oppose that bias. The selection signal thus runs counter to the mutational process rather than arising from it, confirming that the better fit of FMutSel reflects genuine selection on codon usage and not an artefact (Table 2).

Another process that the mutation–selection models cannot by themselves distinguish from selection is GC-biased gene conversion (gBGC), which acts in the same G/C-ward direction as the inferred codon preferences. It is an unlikely driver here, however: recombination among Great Reed Warbler MHC-I haplotypes is rare (2 of 668 transmitted gametes; rate ≈ 0.0030) (*25*), and the duplicated MHC-I genes recombine ectopically in repeat-rich intronic and intergenic sequence rather than in coding exons, biasing breakpoints away from exon 3 (*53*). Moreover, gBGC is indifferent to codon function and cannot explain the stronger match between codon usage and tRNA abundance at conserved sites (Fig. 5) or the association with individual fitness (Fig. 6). I cannot formally partition gBGC from selection in the pooled, multilocus data, but together these observations make it an implausible explanation for the signal.

### An RNA-level arms race

From an ultimate perspective, the release of purifying selection on synonymous variation among certain sites in MHC genes may be driven by an arms race with pathogens (*23*, *56*). Recent investigations uncovered a previously unrecognized antagonistic coevolution of synonymous codon usage between viruses and hosts (*27*). The authors showed that similarity of codon usage between viruses and hosts can be deleterious to the hosts as the expression of virus genes may impede translation of host genes. However, the patterns of synonymous variation observed in the Great Reed Warbler MHC-I exon 3 are inconsistent with that mechanism: two thirds of the sites in the Great Reed Warbler MHC-I exon 3 show evidence of strong purifying selection on codon usage, and this does not suggest a coevolutionary response to avoid similarity with codon usage of viruses. If MHC expression would be affected by viruses matching host codon usage, then reducing the translation of two thirds of the sites would be efficient. It seems unlikely that synonymous variation in the remaining one third of the sites would do much to escape the depletion of tRNAs used by virus genes.

Nonetheless, expression of MHC genes is crucial to vertebrate immunity, and any mechanism that obstructs it would be adaptive to pathogens. It is well established that intracellular parasites such as viruses evade recognition and inhibit host immunity by targeting immune gene expression and protein synthesis, e.g., via molecular interactions with host DNA or mRNA (*28–36*). Such regulation by pathogens is likely to elicit an evolutionary response in the hosts, and I propose the hypothesis that synonymous nucleotide variation in the MHC is engaged in a coevolutionary arms race to escape molecular interactions between MHC mRNA and pathogen molecules, contrasting with the classical protein-level arms race acting on amino acid variation (Fig. 7). This hypothesis is biologically plausible as pathogen targeting of MHC mRNA is established, for example through the Epstein–Barr virus BGLF5 protein that degrades host HLA class I and II mRNAs to evade immune detection (*32*). Due to secondary and tertiary structures of the mRNA, it is likely that not all sites would be equally exposed to molecular interactions with pathogenic RNA or proteins, and this could explain why the synonymous variation appears in spikes (Fig. 1; Fig. 7). Fitness-linked synonymous variation (Fig. 6), concentrated where protein-level selection is absent, is what an RNA-level arms race over MHC expression would produce.

**Fig. 7.**
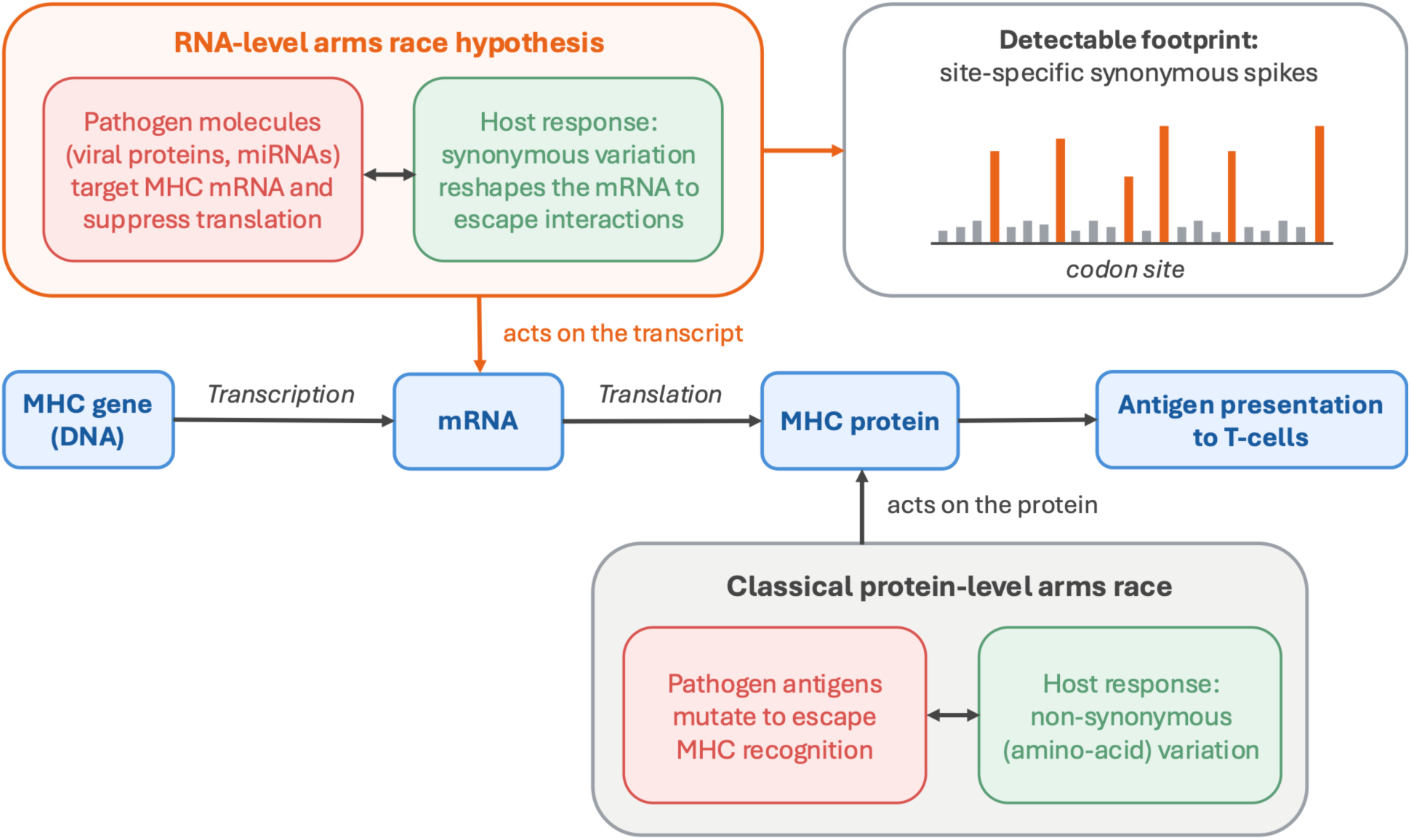
The RNA-level molecular arms-race hypothesis. I propose that MHC evolution is shaped by two host–pathogen arms races: the classical protein-level race, where hosts counter antigen escape through non-synonymous variation, and an RNA-level race, where pathogen molecules (viral proteins, microRNAs) target MHC mRNA to suppress translation and hosts respond through synonymous variation that reshapes the transcript to escape, leaving a site-specific footprint.

Matsushita & Kano-Sueoka interpreted the conspicuous non-random synonymous spikes they described in human HLA-A as a neutral mutational signature of cytosine deamination (76% of synonymous changes were G/C→A/T deamination-type), without testing for selection (*24*). My present results both test and contrast with that interpretation. Deamination is indeed a major contributor to the spikes in the Great Reed Warbler MHC-I (above), but it cannot account for the spikes that fall outside CpG contexts, nor for their association with individual fitness. Where the HLA-A pattern was read as neutral, the Great Reed Warbler MHC-I reveals a layer of selection on synonymous variation beyond the mutational process.

The correlations between synonymous nucleotide variation in the MHC-I and individual survival and fitness in the study population differed between males and females (Fig. 6a, b). This difference in the effects of synonymous variation in the MHC-I is likely associated with regulatory effects of sex hormones on immune responses, that cause males to generally have weaker immune responses and increased risk of pathogen infection compared to females (e.g. (*57–61*)). I have previously proposed that differences between the immune response phenotypes of males and females can drive sexually antagonistic (SA) selection on genes that exert quantitative effects on immune responses, including the MHC, and found evidence for an unresolved sexual conflict over the number of different MHC-I alleles in Great Reed Warblers (*38*, *62*). Additional evidence for sexual conflict on MHC genes recently emerged from a study on a social mammal, the Banded Mongoose *Mungos mungo* (*63*). If the observed sex difference in the effects of synonymous variation in the Great Reed Warbler MHC-I is to be explained by the SA selection hypothesis, then synonymous variation in the MHC-I should fulfill the assumption of exerting quantitative effects on immune responses, cf. (*62*). This is consistent with the hypothesis that synonymous variation serves an adaptive function in evading regulation of MHC expression by pathogens. Polymorphism in MHC genes contributes to quantitative differences in overall antibody production (*64*). Hence, if pathogens regulate MHC expression by targeting MHC mRNA, host mechanisms that evade such regulation would indeed be predicted to exert quantitative effects on immune response phenotypes.

### Implications and future directions

Synonymous variation has been shown to exert multiple effects on the pathway from gene to protein, and the evidence that MHC genes are subject to natural selection on synonymous variation invites reflection over some current practices in MHC research. If pathogens regulate MHC expression by targeting mRNA, then expression of MHC alleles is probably biased in infected cells. The practice of estimating MHC expression by abundance of RNA-derived cDNA needs to be evaluated, as mRNA abundance measured at organism or tissue level may not reflect MHC expression in infected cells. Furthermore, it is common to analyze natural selection on protein sequences by comparing rates of non-synonymous to synonymous variation (dN/dS). This analysis assumes that rates of synonymous variation (dS) can be used as a representation of the background rate of evolution. However, if synonymous variation deviates from the background rate of evolution, such models may fail to provide a meaningful estimate of natural selection on protein sequences (*3*). Yang & Nielsen disputed concerns that natural selection on synonymous nucleotide variation should compromise dN/dS analyses (*4*), but my present results indicate that disregarding selection on synonymous variation leads to substantial overestimation of positive selection in MHC sequences by codeml (Table 2).

More broadly, the inferences available here are limited by the nature of the MHC and our current ability to study it in non-model species. Because exon 3 was sequenced across multiple paralogous MHC-I loci that cannot be individually phased, statistics that assume a single locus, an infinite-sites model, or polarized ancestral-to-derived substitutions (e.g. allele-frequency spectra) cannot be applied without confounding. My conclusions here rest on a convergent set of analyses, each robust to this architecture, because it relies on aggregate codon usage or individual phenotypes rather than single-locus allele frequencies or polarized substitutions: (i) the mutation–selection models, which detect selection on synonymous codon usage directly (Table 2); (ii) the elevated codon usage bias relative to neutral simulations, concentrated at the sites under purifying selection (Fig. 2; Fig. 3); (iii) the correspondence between within-block codon usage and the genomic tRNA pool, likewise strongest at those sites (Fig. 5); and (iv) the individual-level association between synonymous variation and fitness (Fig. 6). Looking ahead, two methodological advances would help resolve the mechanisms that generate and maintain synonymous variation in the MHC. First, null models should establish more fully what synonymous variation looks like under neutrality, incorporating context-dependent mutation (e.g., cytosine deamination), gene conversion, and drift, to provide a baseline against which selection can be inferred. Second, locus-resolved sequencing and phasing would make single-locus allele frequencies and substitution polarization available, allowing site-specific drivers of synonymous variation to be identified directly.

### Conclusions

Here I show that MHC genes are subject to natural selection on synonymous nucleotide variation and that the extent of synonymous variation has direct measurable effects on the Darwinian fitness of individuals. These observations call for a shift in the way that we approach studies of the MHC.

Unraveling the driving forces behind selection on synonymous variation in the MHC is of great importance to our understanding of evolution in this extremely important locus. I have taken a first few steps on the road and shown that most sites in the Great Reed Warbler MHC-I exon 3 are under strong purifying selection on synonymous codon usage, which is associated with the abundance of tRNA isotypes in the genome and likely driven by selection for increased translational efficiency. However, that is only half of the story. Significant spikes of synonymous variation appear in about one third of the sites in the Great Reed Warbler MHC-I exon 3. Crucially, this variation is linked to individual fitness, independently of amino-acid variation and the number of MHC alleles, implicating selection on the synonymous variation itself. I propose that these spikes are best explained in the context of a coevolutionary arms race, where synonymous nucleotide variation serves an adaptive function to escape inhibitory molecular interactions between MHC DNA or mRNA and pathogen molecules. This arms race hypothesis identifies an entirely new dimension of host-pathogen coevolution with direct implications for understanding susceptibility to infectious diseases.

Synonymous variation in the MHC remains largely unexplored, and uncovering the mechanisms by which it exerts its biological effects is central to advancing our understanding of the MHC and host-pathogen coevolution. This study introduces the SynDist function in MHCtools to quantify synonymous nucleotide differences (*65*) and demonstrates a novel approach to test site specific selection on synonymous variation by comparing empirical sequences with synonymous data sets simulated under the assumption of no selection on codon usage. I hope that these methodological advancements will pave the way for future studies on synonymous variation in the MHC.

## Methods

### Data set

This study employed data from previous studies of a wild population of Great Reed Warblers at Lake Kvismare in Sweden (59°10’N, 15°25’E). The data set is based on exhaustive field observations and samples collected in the period 1984–2004 and details on the field methods can be found in (*25*, *38*, *66–71*).

For genetic analyses, I employed the Great Reed Warbler genome assembly (GenBank: ASM2153481v1) (*39*, *40*) in conjunction with a data set of 390 MHC-I exon 3 DNA sequences (*25*, *38*). The MHC-I data set originated from 559 individuals (141 adult males, 131 adult females, and 287 chicks) and the whole genome was sequenced from one of the individuals in the MHC-I data set. The size of the genome assembly was 1.2 Gb in 3,012 scaffolds with scaffold N50 = 21.4 Mb and coverage 45x. The 390 sequences of the MHC-I exon 3 were aligned and trimmed to open reading frame according to conserved residues and sequence motifs (*72*, *73*). The sequences were 87 codons long and contained no gaps. Details on DNA extraction, sequencing, and genotyping of the MHC-I exon 3 can be found in (*25*, *38*). The Great Reed Warbler MHC-I exon 3 sequences are available at GenBank (accession numbers: MH468831–MH469159; MT193762–MT193822).

Fitness analyses were conducted on a subset of samples from the genetic data set for which fitness data has previously been published in (*38*). The fitness data set included lifetime observations of 88 adult males and 100 adult females. These individuals harbored 329 of the 390 MHC-I exon 3 sequences between them, with 6–24 different sequences per individual. Further details can be found in (*38*).

### Selection analyses

I tested 12 substitution models on the Great Reed Warbler MHC-I exon 3 sequences in PhyML version 3.1 (*74*, *75*) using maximum likelihood estimation of nucleotide frequencies and tree topology optimization. Among the 12 models, the generalized time-reversible (GTR) model had the lowest Akaike Information Criterion (AIC) value (Table S22). I used codeml from the PAML software package (*41*, *42*) to test for natural selection on synonymous codon variation among the sequences, using a GTR tree as input. I set codeml to assume one Ω (i.e., dN/dS) ratio for all branches and specified site models M2a (estimating three categories of Ω with one category of sites evolving under positive selection (Ω > 1)) and M8 (estimating a beta distribution for Ω < 1 with an additional category of sites under positive selection (Ω > 1)) (*43*). Within each site model, I employed nested FMutSel vs. FMutSel0 models to test the null hypothesis that variation in codon usage is due to mutation bias alone and not affected by selection acting at synonymous sites (*4*). The log likelihoods of the nested FMutSel and FMutSel0 models were compared using a likelihood ratio test with the formula: 2 × Δln(L) ∼ χ2, with 41 degrees of freedom of the χ2 distribution (reflecting the difference in number of parameters between the models). Sites with Ω > 1 were inferred by Bayes Empirical Bayes analysis for each model (*44*).

To determine whether the selection on codon usage detected by the FMutSel model acts with or against the mutational process, I derived two quantities from the codeml output. First, I recorded the maximum-likelihood nucleotide frequency parameters and transition/transversion ratio (κ) estimated under FMutSel and FMutSel0 (Table S2); under FMutSel these parameters describe the mutational process alone, because selection on codon usage is modelled separately. Second, to summarize the direction of selection within synonymous codon families, thereby avoiding confounding by amino acid composition, I computed, for each degenerate amino acid, the proportion of synonymous codon usage represented by G/C-ending codons, both in the observed sequences and in the equilibrium codon frequencies predicted by FMutSel0 (i.e., under mutation only, with no selection on codon usage; evolver-style frequencies from the codeml output). The two proportions were compared across the 18 degenerate amino acids with a two-tailed paired t-test (Tables S1 and S3). Methionine and tryptophan (single codons) and stop codons were excluded.

### Consensus sequence and prediction of protein folding

I analyzed the frequencies of amino acids in each site among the 390 MHC-I exon 3 sequences and derived a consensus sequence of the amino acids most frequently observed at each site. I employed AlphaFold 3 (*45*) at alphafoldserver.com to predict the folded protein structure from the consensus amino acid sequence (Fig. S1).

### Synonymous genetic variation

All downstream analyses were conducted in R v. 4.5.1 (*76*), except where otherwise specified. Fasta files were handled in R using the package seqinr (*77*). I used the SynDist function in MHCtools v. 1.6 (*65*) to quantify synonymous nucleotide variation among the Great Reed Warbler MHC-I exon 3 sequences. I specified analysis=“codon” to obtain counts of synonymous changes per base and per (codon) site among all pairwise sequence comparisons in the data set. I also used the setting analysis=“dist” to obtain the number of synonymous nucleotide changes in each pairwise sequence comparison in the data set as well as the mean number of synonymous nucleotide changes in pairwise comparisons among the sequences in each individual sample.

I calculated relative frequencies of codon usage as the proportion of counts for a codon out of the total counts for all codons within each synonymous block, i.e., codons encoding the same amino acid. For comparisons between amino acids, I normalized the relative codon frequencies to 1 by multiplying each frequency with the number of synonymous codons in each block, thereby generating the measure “relative synonymous codon usage” (RSCU), cf. (*10*, *78*). I quantified codon usage bias (CUB) for each amino acid as the variance of RSCU within each synonymous block, following the rationale that a bias towards certain codons will skew the values of RSCU within each synonymous block and increase the variance. Thirty sites with an observed frequency of synonymous variation > 0.03 were classified as showing relaxed purifying selection on codon usage; the remaining 57 sites were classified as evolving under strong purifying selection.

### Data simulations

If one assumes that an observed sequence alignment is a snapshot of a Markov process of synonymous codon substitutions that has run for infinitely long time, the probability of observing a certain codon can be regarded as a function of the mutation bias among nucleotides and selection on codon usage, following (*4*). Hence, under the null hypothesis of no selection on codon usage, the probability of observing a certain codon depends only on mutation bias, and thus regarding the process of synonymous codon substitution as a Markov process enables simulation of nucleotide sequences that reflect the null hypothesis using mutation bias parameters alone. I employed custom scripts to perform simulations of MHC-I exon 3 sequences reflecting the null hypothesis that variation in codon usage is not affected by selection acting at synonymous sites. I first used the base frequencies by nucleotide position (3 × 4 table) derived from the models in codeml (Table S23) to compute a table of relative probabilities of codon usage under neutrality, i.e., reflecting the scenario where the probabilities of observing synonymous codon variants depend only on the mutation bias, cf. (*4*) (Table S24). I then ran 1,000 simulations of the data set of 390 MHC-I exon 3 sequences, where for each site in each sequence, a codon was randomly sampled among the synonymous codons encoding the observed amino acid. In the random sampling step, the relative probabilities of synonymous codons encoding the observed amino acid in the computed table of probabilities of codon usage under neutrality (Table S24) were used as weights. The simulated codons were concatenated into nucleotide sequences and collated in 1,000 fasta files, each containing 390 simulated DNA sequences synonymous to the empirical Great Reed Warbler MHC-I exon 3 sequences.

I then used the SynDist function in MHCtools v. 1.6 to quantify the synonymous genetic variation among the 390 sequences in each of the 1,000 simulated data sets. I specified analysis=“codon” to obtain counts of synonymous changes per base and per (codon) site among all pairwise sequence comparisons in the data set. From the results, I extracted the proportion of pairwise sequence comparisons that harbor synonymous substitutions for each site. In addition, I calculated relative frequencies of codon usage, RSCU, and codon usage bias as described above. Finally, I calculated means and 95% confidence intervals among the 1,000 simulated data sets and calculated p-values as the proportion of observations in the simulated data sets more extreme than the value observed in the empirical data set.

### CpG context of synonymous variation

To test whether cytosine deamination could account for the site-specific synonymous variation, I examined the sequence context of the 87 codon sites. From a majority-rule consensus of the 390 MHC-I exon 3 sequences, I identified CpG dinucleotides and mapped them to codon positions, recording for each site whether its third position (i) was a cytosine or guanine, the deamination substrate, and (ii) fell within a CpG dinucleotide (including CpGs spanning the codon boundary). I compared the sites under strong purifying selection (n = 57) and relaxed sites (n = 30) against each context with Fisher’s exact tests.

### Abundance of tRNA isotypes

I used tRNAscan-SE v. 2.0.12 (*46*) to scan the Great Reed Warbler genome assembly for tRNA genes. I ran tRNAscan-SE with the options ‘--mt vert’ to identify potential mitochondrial origin of detected tRNAs and ‘--detail’ to obtain isotype-specific model classification results. tRNAscan-SE predicted 815 tRNAs in the first pass, 571 of which were confirmed by Infernal (second pass). Among the 571 detected tRNAs, 325 decoded standard amino acids, 1 was a selenocysteine tRNA, 7 had undetermined isotypes, 30 had mismatch isotypes, and 208 were pseudogenes. I used only the 325 confirmed cytosolic standard amino acid tRNAs in further analyses.

I calculated relative frequencies of tRNA isotypes as the proportion of counts for an isotype out of the total counts for isotypes for the same amino acid. For comparisons between amino acids, I normalized the relative tRNA isotype frequencies by multiplying each frequency with the number of synonymous codons in each block.

To test whether synonymous codon usage was affected by tRNA isotype availability, I calculated Pearson correlation coefficients between RSCU and normalized relative frequencies of tRNA isotypes within each synonymous block. The analysis included only amino acids that were matched by more than one tRNA isotype in the genome and excluded codons that were not matched by a cognate tRNA isotype in the genome. The Pearson correlations were calculated using RSCU measured among all 87 sites of the MHC-I sequences as well as RSCU measured among subsets of sites where synonymous variation was under strong purifying selection (n = 57) or where purifying selection was relaxed (n = 30). Pearson correlations were also calculated using RSCU for each of the 1,000 data sets simulated under the assumption of no selection on synonymous codon usage, allowing derivation of 95% confidence intervals and p-values for the correlations between normalized relative frequencies of tRNA isotypes and RSCU in the empirical data set.

### Fitness effects

As estimates of individual Darwinian fitness, I used *lifetime number of fledged offspring*, defined as the number of fledged offspring over an individual’s lifetime, and *offspring fledging success*, defined as the lifetime number of fledged offspring corrected for life span. Previous studies have shown that Great Reed Warblers that failed their first breeding attempt were unlikely to be observed in the study area in following years (*38*, *67*, *79*). Estimates of life span and lifetime reproductive success are therefore not reliable for unsuccessful first-time breeders and I excluded 17 individuals that failed to rear any offspring from analyses of fitness effects, resulting in a data set of 77 males and 96 females.

I used generalized linear models to test the effects of synonymous genetic variation in the Great Reed Warbler MHC-I exon 3 on *life span, lifetime number of fledged offspring*, and *offspring fledging success* following the suggested model design in (*80*). As both *life span* and *lifetime number of fledged offspring* follow negative binomial distributions, the generalized linear models were run with negative binomial errors using the package MASS in R (*81*). The aggregation parameters of the negative binomial distributions were estimated by maximum likelihood and specified in the model formulae (*82*). Effects on *offspring fledging success* were modelled by including *life span* as covariate in a generalized linear model with *lifetime number of fledged offspring* as dependent variable.

The purpose of the models was to test the effects of the mean number of synonymous nucleotide changes in pairwise comparisons among the sequences in each individual sample. The full models included the total number of different MHC-I sequences per individual as covariate, sex as fixed factor, and the interactions ‘total number of different MHC-I sequences × sex’ and ‘mean number of synonymous nucleotide changes × sex’. The step function in R was employed for model simplification and simulateResiduals and testDispersion from the DHARMa package (*83*) were used for model diagnostics. The final models on *life span* and *lifetime number of fledged offspring* showed significant or nearly significant interactions between the mean number of synonymous nucleotide changes and sex, and the models were therefore also run independently for each sex.

To ascertain that the observed effects of synonymous variation were not influenced by natural selection on amino acid sequences, I calculated the mean proportion of amino acid changes between pairs of sequences in each individual (i.e., amino acid p-distance) using the DistCalc function in MHCtools v. 1.6 (*65*). I repeated the generalized linear models as above including amino acid p-distance as covariate and the interaction ‘amino acid p-distance × sex’ in the full models. In addition, I repeated the models as described above, but replacing the mean number of synonymous nucleotide changes with amino acid p-distance as independent variable.

## Acknowledgements

I wish to thank my partner Tianhao Zhao for patience and support. I also wish to thank the Great Reed Warbler project at Lund University and everyone who has contributed to the field work.

In memory of Bengt Olle Bengtsson (1946–2025).

## Author contributions

This study was conceived, designed and carried out by J.R. MHCtools is developed and maintained by J.R. The data simulation approach to test site specific selection on synonymous variation was conceived and developed by J.R.

## Data and materials availability

MHCtools v. 1.6 including user manual and documentation is available at CRAN: https://cran.r-project.org/package=MHCtools. The data sets are available at the Dryad repository: https://datadryad.org/dataset/doi:10.5061/dryad.b321hf1 and the Zenodo repository: https://doi.org/10.5281/zenodo.3716048. The great reed warbler genome assembly and MHC-I exon 3 sequences are available at GenBank: https://ncbi.nlm.nih.gov (accession numbers: MH468831–MH469159; MT193762–MT193822; genome assembly: ASM2153481v1).

