## Supplementary tables and figures for "Natural selection on synonymous genetic variation in the major histocompatibility complex"

**Table of contents**


| Page(s) | |
|---|---|
| 2 | Table S1, Table S2 |
| 3–4 | Table S3 |
| 5 | Table S4, Table S5 |
| 6 | Table S6, Table S7 |
| 7–9 | Table S8 |
| 9 | Table S9, Table S10 |
| 10 | Table S11 |
| 11 | Table S12 |
| 12–14 | Table S13 |
| 15 | Table S14 |
| 16 | Table S15 |
| 17 | Table S16, Table S17 |
| 18 | Table S18, Table S19 |
| 19 | Table S20, Table S21 |
| 20 | Table S22, Table S23 |
| 21 | Table S24 |
| 22 | Fig. S1 |
| 23 | Fig. S2 |
| 24 | Fig. S3 |

**Table S1.** *Within each synonymous block, the proportion of codon usage in G/C-ending codons among the 390 MHC-I sequences and corresponding mutation-only expectations from the M8 mutation–selection models in codeml. Observed usage is shifted toward G/C-ending codons in 16 of 18 blocks (mean 70.1% vs. 40.5%; mean difference +29.6 percentage points), showing that selection on codon usage acts in a net G/C-ward direction, opposite to the A/T-biased mutational process. Because the FMutSel0 equilibrium already carries an inflated G/C mutation frequency, this comparison is conservative. %GC3 specifies the percentage of within-block codon usage in codons ending in G or C.*

| Amino acid | codons (k) | N (obs) | Observed %GC3 | Mutation-only %GC3 | Difference |
|---|---|---|---|---|---|
| Ala | 4 | 1788 | 35.5 | 41.3 | -5.8 |
| Arg | 6 | 4111 | 62.5 | 42.5 | +20.0 |
| Asn | 2 | 534 | 6.2 | 38.3 | -32.1 |
| Asp | 2 | 2039 | 70.0 | 38.3 | +31.7 |
| Cys | 2 | 786 | 49.9 | 38.3 | +11.6 |
| Gln | 2 | 927 | 98.5 | 45.8 | +52.6 |
| Glu | 2 | 3731 | 63.8 | 45.8 | +17.9 |
| Gly | 4 | 3413 | 77.6 | 41.3 | +36.3 |
| His | 2 | 1102 | 57.6 | 38.3 | +19.3 |
| Ile | 3 | 1220 | 95.9 | 28.3 | +67.6 |
| Leu | 6 | 2891 | 96.3 | 43.1 | +53.2 |
| Lys | 2 | 1172 | 77.9 | 45.8 | +32.1 |
| Phe | 2 | 1336 | 73.8 | 38.3 | +35.5 |
| Pro | 4 | 523 | 71.9 | 41.3 | +30.6 |
| Ser | 6 | 2324 | 81.1 | 40.6 | +40.4 |
| Thr | 4 | 1173 | 80.8 | 41.3 | +39.5 |
| Tyr | 2 | 1843 | 86.9 | 38.3 | +48.6 |
| Val | 4 | 1830 | 75.5 | 41.3 | +34.2 |
| **Mean/test** | | | **70.1** | **40.5** | **+29.6** |

**Table S2.** *Maximum-likelihood nucleotide frequency parameters and κ under the mutation–selection model (M8 FMutSel) and the mutation-only null model (M8 FMutSel0) in codeml. Under FMutSel the mutational process is strongly A/T-biased (G+C = 0.267), consistent with cytosine deamination; FMutSel0, lacking selection on codon usage, instead infers a higher G/C frequency (0.413) to accommodate the G/C-rich observed usage.*

| Model | π(T) | π(C) | π(A) | π(G) | G+C | κ |
|---|---|---|---|---|---|---|
| FMutSel | 0.516 | 0.165 | 0.217 | 0.102 | 0.267 | 2.46 |
| FMutSel0 | 0.374 | 0.232 | 0.214 | 0.181 | 0.413 | 2.46 |

**Table S3.** *Observed codon usage counts, observed relative synonymous codon usage (RSCU), and the mutation-only RSCU expected from the M8 FMutSel0 equilibrium codon frequencies, per codon. Mutation-only RSCU is computed from the FMutSel0 equilibrium (evolver-style) codon frequencies. • marks codons ending in G or C.*

| Amino acid | Codon | 3' G/C | Observed count | Observed RSCU | Mutation-only RSCU |
|---|---|---|---|---|---|
| Ala | GCT | | 1126 | 2.519 | 1.495 |
| Ala | GCC | • | 362 | 0.810 | 0.928 |
| Ala | GCA | | 27 | 0.060 | 0.854 |
| Ala | GCG | • | 273 | 0.611 | 0.723 |
| Arg | CGT | | 434 | 0.633 | 1.645 |
| Arg | CGC | • | 722 | 1.054 | 1.021 |
| Arg | CGA | | 256 | 0.374 | 0.940 |
| Arg | CGG | • | 901 | 1.315 | 0.796 |
| Arg | AGA | | 853 | 1.245 | 0.866 |
| Arg | AGG | • | 945 | 1.379 | 0.733 |
| Asn | AAT | | 501 | 1.876 | 1.234 |
| Asn | AAC | • | 33 | 0.124 | 0.766 |
| Asp | GAT | | 611 | 0.599 | 1.234 |
| Asp | GAC | • | 1428 | 1.401 | 0.766 |
| Cys | TGT | | 394 | 1.003 | 1.234 |
| Cys | TGC | • | 392 | 0.997 | 0.766 |
| Gln | CAA | | 14 | 0.030 | 1.083 |
| Gln | CAG | • | 913 | 1.970 | 0.917 |
| Glu | GAA | | 1352 | 0.725 | 1.083 |
| Glu | GAG | • | 2379 | 1.275 | 0.917 |
| Gly | GGT | | 104 | 0.122 | 1.495 |
| Gly | GGC | • | 775 | 0.908 | 0.928 |
| Gly | GGA | | 660 | 0.774 | 0.854 |
| Gly | GGG | • | 1874 | 2.196 | 0.723 |
| His | CAT | | 467 | 0.848 | 1.234 |
| His | CAC | • | 635 | 1.152 | 0.766 |
| Ile | ATT | | 15 | 0.037 | 1.368 |
| Ile | ATC | • | 1170 | 2.877 | 0.849 |
| Ile | ATA | | 35 | 0.086 | 0.782 |
| Leu | TTA | | 2 | 0.004 | 1.262 |
| Leu | TTG | • | 480 | 0.996 | 1.069 |
| Leu | CTT | | 96 | 0.199 | 1.371 |
| Leu | CTC | • | 756 | 1.569 | 0.851 |
| Leu | CTA | | 9 | 0.019 | 0.784 |
| Leu | CTG | • | 1548 | 3.213 | 0.663 |
| Lys | AAA | | 259 | 0.442 | 1.083 |
| Lys | AAG | • | 913 | 1.558 | 0.917 |
| Phe | TTT | | 350 | 0.524 | 1.234 |
| Phe | TTC | • | 986 | 1.476 | 0.766 |

| Amino acid | Codon | 3' G/C | Observed count | Observed RSCU | Mutation-only RSCU |
|---|---|---|---|---|---|
| Pro | CCT | | 120 | 0.918 | 1.495 |
| Pro | CCC | • | 3 | 0.023 | 0.928 |
| Pro | CCA | | 27 | 0.207 | 0.854 |
| Pro | CCG | • | 373 | 2.853 | 0.723 |
| Ser | TCT | | 263 | 0.679 | 1.766 |
| Ser | TCC | • | 1210 | 3.124 | 1.096 |
| Ser | TCA | | 1 | 0.003 | 1.009 |
| Ser | TCG | • | 0 | 0.000 | 0.854 |
| Ser | AGT | | 176 | 0.454 | 0.787 |
| Ser | AGC | • | 674 | 1.740 | 0.488 |
| Thr | ACT | | 52 | 0.177 | 1.495 |
| Thr | ACC | • | 723 | 2.465 | 0.928 |
| Thr | ACA | | 173 | 0.590 | 0.854 |
| Thr | ACG | • | 225 | 0.767 | 0.723 |
| Tyr | TAT | | 242 | 0.263 | 1.234 |
| Tyr | TAC | • | 1601 | 1.737 | 0.766 |
| Val | GTT | | 448 | 0.979 | 1.495 |
| Val | GTC | • | 746 | 1.631 | 0.928 |
| Val | GTA | | 1 | 0.002 | 0.854 |
| Val | GTG | • | 635 | 1.388 | 0.723 |

**Table S4** *Summary of the M2a mutation-selection vs. null models in codeml. ln(L) specifies the log likelihood of the models, Ω > 1 specifies the proportion of sites that were estimated to be under positive selection for amino acid change, and Ω est indicates the value of Ω estimated for that category of sites. P-value was calculated by likelihood ratio test of the nested FMutSel vs. FMutSel0 models.*

| Site model | Codon model | Hypothesis | ln(L) | Ω > 1 | Ω est | $P(H_0)$ |
|---|---|---|---|---|---|---|
| M2a | FMutSel0 | $H_0$: No selection on synonymous variation | -7521.0 | 16.8% | 4.42 | 3.16e-14 |
| | FMutSel | $H_1$: Selection on synonymous variation | -7446.4 | 15.2% | 3.45 | |

**Table S5** *Positively selected sites inferred by Bayes Empirical Bayes analysis in the M8 mutation-selection model (FMutSel). Note that the p-value for site 55 is not significant.*

| Site | P (Ω > 1) | Post mean Ω | S.E. for Ω |
|---|---|---|---|
| 1 | 1.000 | 3.500 | 0.013 |
| 3 | 1.000 | 3.500 | 0.013 |
| 5 | 1.000 | 3.500 | 0.013 |
| 21 | 1.000 | 3.500 | 0.013 |
| 32 | 0.974 | 3.434 | 0.406 |
| 34 | 0.998 | 3.494 | 0.119 |
| 44 | 1.000 | 3.500 | 0.016 |
| 55 | 0.930 | 3.320 | 0.656 |
| 58 | 1.000 | 3.500 | 0.013 |
| 61 | 1.000 | 3.500 | 0.013 |
| 62 | 1.000 | 3.500 | 0.013 |
| 69 | 1.000 | 3.500 | 0.013 |
| 79 | 0.968 | 3.418 | 0.452 |

**Table S6** *Summary of the codon variation in site 61 among the 390 MHC-I sequences. The synonymous variation in the first nucleotide position can be ascribed to the 6-fold degenerate amino acid arginine, which appears in 234 of the 390 sequences.*

| Amino acid | | Codon | Count |
|---|---|---|---|
| Ser | S | AGT | 60 |
| His | H | CAC | 20 |
| | | CAT | 46 |
| Arg | R | AGG | 123 |
| | | CGT | 48 |
| | | CGG | 13 |
| | | AGA | 50 |
| Trp | W | TGG | 1 |
| Gly | G | GGT | 3 |
| Tyr | Y | TAT | 14 |
| Lys | K | AAA | 11 |
| Thr | T | ACT | 1 |

**Table S7** *Location of encoded amino acids in bridge domains, β-sheets, or α-helices of the folded protein structure of the consensus amino acid sequence derived from the 390 Great Reed Warbler MHC-I exon 3 sequences as predicted by AlphaFold 3 (Fig. S1).*

| Site | Domain type |
|---|---|
| 1 | bridge |
| 2-9 | β-sheet |
| 10-14 | bridge |
| 15-23 | β-sheet |
| 24-25 | bridge |
| 26-32 | β-sheet |
| 33-36 | bridge |
| 37-40 | β-sheet |
| 41-42 | bridge |
| 43-55 | α-helix |
| 56 | bridge |
| 57-67 | α-helix |
| 68 | bridge |
| 69-85 | α-helix |
| 86-87 | bridge |

**Table S8** *Frequencies of synonymous changes for each site among the 390 Great Reed Warbler MHC-I exon 3 sequences (Freq. syn. obs.) and the mean among the sequences in the 1,000 data sets simulated under the assumption of no selection on synonymous variation (Mean sim.). C.I. lower and C.I. upper indicate the boundaries of 95% confidence intervals for the synonymous variation in the simulated data sets, and the p-values indicate the proportion of observations in the simulated data sets more extreme than the value observed in the empirical data set. Sign indicates whether the observed empirical value was lower or higher than the mean among the simulated data sets.*

| Site | Freq. syn. obs. | Mean sim. | C.I. lower | C.I. upper | P-value | Sign |
|---|---|---|---|---|---|---|
| 1 | 0.048 | 0.122 | 0.118 | 0.125 | 0.000 | - |
| 2 | 0.000 | 0.631 | 0.617 | 0.642 | 0.000 | - |
| 3 | 0.127 | 0.448 | 0.438 | 0.457 | 0.000 | - |
| 4 | 0.005 | 0.565 | 0.545 | 0.581 | 0.000 | - |
| 5 | 0.119 | 0.169 | 0.155 | 0.179 | 0.000 | - |
| 6 | 0.000 | 0.676 | 0.654 | 0.696 | 0.000 | - |
| 7 | 0.000 | 0.425 | 0.382 | 0.458 | 0.000 | - |
| 8 | 0.002 | 0.206 | 0.178 | 0.232 | 0.000 | - |
| 9 | 0.000 | 0.741 | 0.726 | 0.754 | 0.000 | - |
| 10 | 0.020 | 0.788 | 0.773 | 0.802 | 0.000 | - |
| 11 | 0.220 | 0.772 | 0.751 | 0.792 | 0.000 | - |
| 12 | 0.221 | 0.415 | 0.377 | 0.447 | 0.000 | - |
| 13 | 0.000 | 0.670 | 0.649 | 0.688 | 0.000 | - |
| 14 | 0.392 | 0.757 | 0.735 | 0.774 | 0.000 | - |
| 15 | 0.004 | 0.457 | 0.440 | 0.471 | 0.000 | - |
| 16 | 0.000 | 0.710 | 0.696 | 0.722 | 0.000 | - |
| 17 | 0.380 | 0.695 | 0.673 | 0.715 | 0.000 | - |
| 18 | 0.152 | 0.372 | 0.361 | 0.381 | 0.000 | - |
| 19 | 0.000 | 0.154 | 0.137 | 0.168 | 0.000 | - |
| 20 | 0.000 | 0.760 | 0.745 | 0.772 | 0.000 | - |
| 21 | 0.018 | 0.086 | 0.078 | 0.093 | 0.000 | - |
| 22 | 0.020 | 0.687 | 0.665 | 0.707 | 0.000 | - |
| 23 | 0.205 | 0.426 | 0.387 | 0.457 | 0.000 | - |
| 24 | 0.115 | 0.387 | 0.352 | 0.418 | 0.000 | - |
| 25 | 0.019 | 0.594 | 0.575 | 0.611 | 0.000 | - |
| 26 | 0.017 | 0.540 | 0.529 | 0.550 | 0.000 | - |
| 27 | 0.000 | 0.419 | 0.383 | 0.450 | 0.000 | - |
| 28 | 0.000 | 0.421 | 0.385 | 0.455 | 0.000 | - |
| 29 | 0.000 | 0.573 | 0.535 | 0.605 | 0.000 | - |
| 30 | 0.000 | 0.760 | 0.740 | 0.778 | 0.000 | - |

| | | | | | | |
|---|---|---|---|---|---|---|
| 31 | 0.470 | 0.399 | 0.362 | 0.432 | 0.000 | + |
| 32 | 0.000 | 0.200 | 0.181 | 0.219 | 0.000 | - |
| 33 | 0.005 | 0.792 | 0.778 | 0.806 | 0.000 | - |
| 34 | 0.002 | 0.235 | 0.213 | 0.257 | 0.000 | - |
| 35 | 0.074 | 0.769 | 0.748 | 0.788 | 0.000 | - |
| 36 | 0.000 | 0.582 | 0.563 | 0.597 | 0.000 | - |
| 37 | 0.000 | 0.780 | 0.765 | 0.793 | 0.000 | - |
| 38 | 0.264 | 0.423 | 0.381 | 0.455 | 0.000 | - |
| 39 | 0.000 | 0.534 | 0.517 | 0.549 | 0.000 | - |
| 40 | 0.074 | 0.376 | 0.364 | 0.386 | 0.000 | - |
| 41 | 0.114 | 0.677 | 0.656 | 0.696 | 0.000 | - |
| 42 | 0.000 | 0.423 | 0.383 | 0.455 | 0.000 | - |
| 43 | 0.005 | 0.776 | 0.755 | 0.794 | 0.000 | - |
| 44 | 0.000 | 0.484 | 0.468 | 0.499 | 0.000 | - |
| 45 | 0.005 | 0.694 | 0.673 | 0.713 | 0.000 | - |
| 46 | 0.000 | 0.348 | 0.302 | 0.395 | 0.000 | - |
| 47 | 0.000 | 0.576 | 0.539 | 0.609 | 0.000 | - |
| 48 | 0.010 | 0.614 | 0.594 | 0.631 | 0.000 | - |
| 49 | 0.000 | 0.792 | 0.777 | 0.805 | 0.000 | - |
| 50 | 0.000 | 0.792 | 0.776 | 0.805 | 0.000 | - |
| 51 | 0.000 | 0.619 | 0.605 | 0.630 | 0.000 | - |
| 52 | 0.000 | 0.000 | 0.000 | 0.001 | 0.000 | - |
| 53 | 0.160 | 0.350 | 0.302 | 0.390 | 0.000 | - |
| 54 | 0.040 | 0.338 | 0.306 | 0.365 | 0.000 | - |
| 55 | 0.000 | 0.262 | 0.225 | 0.296 | 0.000 | - |
| 56 | 0.005 | 0.695 | 0.674 | 0.714 | 0.000 | - |
| 57 | 0.062 | 0.318 | 0.304 | 0.332 | 0.000 | - |
| 58 | 0.000 | 0.172 | 0.161 | 0.182 | 0.000 | - |
| 59 | 0.000 | 0.694 | 0.673 | 0.713 | 0.000 | - |
| 60 | 0.005 | 0.349 | 0.302 | 0.392 | 0.000 | - |
| 61 | 0.241 | 0.316 | 0.308 | 0.323 | 0.000 | - |
| 62 | 0.000 | 0.070 | 0.062 | 0.079 | 0.000 | - |
| 63 | 0.426 | 0.619 | 0.599 | 0.636 | 0.000 | - |
| 64 | 0.000 | 0.425 | 0.387 | 0.460 | 0.000 | - |
| 65 | 0.000 | 0.423 | 0.386 | 0.454 | 0.000 | - |
| 66 | 0.154 | 0.784 | 0.770 | 0.798 | 0.000 | - |
| 67 | 0.005 | 0.320 | 0.275 | 0.364 | 0.000 | - |
| 68 | 0.043 | 0.386 | 0.351 | 0.417 | 0.000 | - |
| 69 | 0.034 | 0.196 | 0.174 | 0.218 | 0.000 | - |

| | | | | | | |
|---|---|---|---|---|---|---|
| 70 | 0.000 | 0.422 | 0.386 | 0.454 | 0.000 | - |
| 71 | 0.508 | 0.691 | 0.670 | 0.711 | 0.000 | - |
| 72 | 0.088 | 0.348 | 0.299 | 0.390 | 0.000 | - |
| 73 | 0.000 | 0.006 | 0.006 | 0.007 | 0.000 | - |
| 74 | 0.075 | 0.792 | 0.776 | 0.805 | 0.000 | - |
| 75 | 0.000 | 0.331 | 0.287 | 0.374 | 0.000 | - |
| 76 | 0.000 | 0.287 | 0.268 | 0.302 | 0.000 | - |
| 77 | 0.096 | 0.212 | 0.192 | 0.228 | 0.000 | - |
| 78 | 0.187 | 0.684 | 0.663 | 0.703 | 0.000 | - |
| 79 | 0.000 | 0.381 | 0.371 | 0.389 | 0.000 | - |
| 80 | 0.356 | 0.425 | 0.385 | 0.458 | 0.002 | - |
| 81 | 0.000 | 0.694 | 0.671 | 0.713 | 0.000 | - |
| 82 | 0.000 | 0.310 | 0.267 | 0.352 | 0.000 | - |
| 83 | 0.000 | 0.344 | 0.300 | 0.389 | 0.000 | - |
| 84 | 0.000 | 0.349 | 0.302 | 0.392 | 0.000 | - |
| 85 | 0.045 | 0.788 | 0.775 | 0.801 | 0.000 | - |
| 86 | 0.010 | 0.346 | 0.301 | 0.391 | 0.000 | - |
| 87 | 0.000 | 0.740 | 0.726 | 0.754 | 0.000 | - |

**Table S9** *Summary of the codon variation in site 31 among the 390 MHC-I sequences. The synonymous variation can be ascribed to the 2-fold degenerate amino acid phenylalanine, which appears in 378 of the 390 sequences.*

| Amino acid | | Codon | Count |
|---|---|---|---|
| Phe | F | TTT | 195 |
| | | TTC | 183 |
| Leu | L | CTT | 12 |

**Table S10** *Counts of C/G and CpG at 3rd codon position in the 57 sites where codon usage was under strong purifying selection and the 30 sites where purifying selection was relaxed.*

| Site class | CG3 | | CpG3 | |
|---|---|---|---|---|
| | FALSE | TRUE | FALSE | TRUE |
| Purifying sel. | 14 | 43 | 56 | 1 |
| Relaxed sel. | 8 | 22 | 15 | 15 |

**Table S11** *Codon usage bias (CUB) measured as the variance of RSCU among the 390 empirical MHC-I exon 3 sequences and the mean among the 1,000 data sets simulated under the assumption of no selection on synonymous variation (Mean sim.). C.I. lower and C.I. upper indicate the boundaries of 95% confidence intervals for CUB in the simulated data sets, and the p-values indicate the proportion of observations in the simulated data sets more extreme than the value observed in the empirical data set.*

| Amino acid | | CUB | | | | |
|---|---|---|---|---|---|---|
| | | Empirical | Mean sim. | C.I. lower | C.I. upper | P-value |
| Phe | F | 0.45 | 0.30 | 0.23 | 0.39 | 0.001 |
| Leu | L | 1.56 | 0.30 | 0.26 | 0.34 | 0 |
| Ile | I | 2.64 | 0.41 | 0.33 | 0.50 | 0 |
| Met | M | NA | NA | NA | NA | NA |
| Val | V | 0.51 | 0.30 | 0.25 | 0.35 | 0 |
| Ser | S | 1.49 | 0.41 | 0.35 | 0.48 | 0 |
| Pro | P | 1.67 | 0.30 | 0.21 | 0.40 | 0 |
| Thr | T | 1.02 | 0.30 | 0.24 | 0.37 | 0 |
| Ala | A | 1.13 | 0.30 | 0.25 | 0.35 | 0 |
| Tyr | Y | 1.09 | 0.30 | 0.23 | 0.36 | 0 |
| His | H | 0.05 | 0.30 | 0.22 | 0.38 | 0 |
| Gln | Q | 1.88 | 0.61 | 0.49 | 0.73 | 0 |
| Asn | N | 1.54 | 0.30 | 0.18 | 0.42 | 0 |
| Lys | K | 0.62 | 0.61 | 0.51 | 0.71 | 0.38 |
| Asp | D | 0.32 | 0.30 | 0.24 | 0.37 | 0.27 |
| Glu | E | 0.15 | 0.61 | 0.55 | 0.67 | 0 |
| Cys | C | 0.00 | 0.30 | 0.21 | 0.42 | 0 |
| Trp | W | NA | NA | NA | NA | NA |
| Arg | R | 0.17 | 0.30 | 0.27 | 0.33 | 0 |
| Gly | G | 0.75 | 0.30 | 0.26 | 0.34 | 0 |

**Table S12** *Codon usage bias (CUB) measured as the variance of RSCU among the 390 MHC-I exon 3 sequences in the 57 sites where codon usage was under strong purifying selection and the 30 sites where purifying selection was relaxed, and the difference between them.*

| Amino acid | | CUB | | |
|---|---|---|---|---|
| | | Purifying sel. | Relaxed sel. | Difference |
| Phe | F | 0.90 | 0.22 | -0.68 |
| Leu | L | 1.34 | 2.32 | 0.98 |
| Ile | I | 3.00 | 1.78 | -1.22 |
| Met | M | NA | NA | NA |
| Val | V | 0.53 | 1.47 | 0.94 |
| Ser | S | 2.31 | 1.42 | -0.90 |
| Pro | P | 3.57 | 1.61 | -1.96 |
| Thr | T | 3.27 | 0.51 | -2.77 |
| Ala | A | 3.72 | 1.04 | -2.69 |
| Tyr | Y | 1.98 | 0.71 | -1.26 |
| His | H | 0.86 | 0.02 | -0.84 |
| Gln | Q | 1.99 | 0.50 | -1.49 |
| Asn | N | 1.80 | 0.89 | -0.91 |
| Lys | K | 0.81 | 0.30 | -0.52 |
| Asp | D | 0.05 | 1.29 | 1.23 |
| Glu | E | 0.58 | 0.37 | -0.21 |
| Cys | C | 0.00 | 0.00 | 0.00 |
| Trp | W | NA | NA | NA |
| Arg | R | 0.32 | 0.69 | 0.37 |
| Gly | G | 1.15 | 1.92 | 0.76 |

**Table S13** *Counts and normalized relative frequencies of tRNA isotypes in the Great Reed Warbler genome and codon counts and RSCU among the 390 MHC-I exon 3 sequences. The four right-most columns show the counts and relative frequencies of codons among the 57 sites where codon usage was under strong purifying selection and the 30 sites where purifying selection was relaxed.*

| Amino acid | | Codon | Anticodon | tRNA | | All sites | | Purifying sel. | | Relaxed sel. | |
|---|---|---|---|---|---|---|---|---|---|---|---|
| | | | | Counts | Norm.Rel.Freq. | Counts | RSCU | Counts | RSCU | Counts | RSCU |
| Phe | F | TTT | AAA | 0 | 0.000 | 350 | 0.524 | 94 | 0.330 | 256 | 0.668 |
| Phe | F | TTC | GAA | 7 | 2.000 | 986 | 1.476 | 475 | 1.670 | 511 | 1.332 |
| Leu | L | TTA | TAA | 1 | 0.375 | 2 | 0.004 | 1 | 0.004 | 1 | 0.005 |
| Leu | L | TTG | CAA | 2 | 0.750 | 480 | 0.996 | 470 | 1.658 | 10 | 0.050 |
| Leu | L | CTT | AAG | 6 | 2.250 | 96 | 0.199 | 41 | 0.145 | 55 | 0.277 |
| Leu | L | CTC | GAG | 0 | 0.000 | 756 | 1.569 | 381 | 1.344 | 375 | 1.891 |
| Leu | L | CTA | TAG | 2 | 0.750 | 9 | 0.019 | 0 | 0.000 | 9 | 0.045 |
| Leu | L | CTG | CAG | 5 | 1.875 | 1548 | 3.213 | 808 | 2.850 | 740 | 3.731 |
| Ile | I | ATT | AAT | 46 | 2.300 | 15 | 0.037 | 0 | 0.000 | 15 | 0.139 |
| Ile | I | ATC | GAT | 12 | 0.600 | 1170 | 2.877 | 896 | 3.000 | 274 | 2.537 |
| Ile | I | ATA | TAT | 2 | 0.100 | 35 | 0.086 | 0 | 0.000 | 35 | 0.324 |
| Met | M | ATG | CAT | 15 | 1.000 | 116 | 1.000 | 93 | 1.000 | 23 | 1.000 |
| Val | V | GTT | AAC | 4 | 1.600 | 448 | 0.979 | 370 | 1.237 | 78 | 0.492 |
| Val | V | GTC | GAC | 1 | 0.400 | 746 | 1.631 | 309 | 1.033 | 437 | 2.757 |
| Val | V | GTA | TAC | 1 | 0.400 | 1 | 0.002 | 0 | 0.000 | 1 | 0.006 |
| Val | V | GTG | CAC | 4 | 1.600 | 635 | 1.388 | 517 | 1.729 | 118 | 0.744 |
| Ser | S | TCT | AGA | 12 | 2.769 | 263 | 0.679 | 0 | 0.000 | 263 | 1.031 |
| Ser | S | TCC | GGA | 0 | 0.000 | 1210 | 3.124 | 387 | 2.928 | 823 | 3.225 |
| Ser | S | TCA | TGA | 3 | 0.692 | 1 | 0.003 | 1 | 0.008 | 0 | 0.000 |
| Ser | S | TCG | CGA | 6 | 1.385 | 0 | 0.000 | 0 | 0.000 | 0 | 0.000 |
| Pro | P | CCT | AGG | 7 | 2.154 | 120 | 0.918 | 0 | 0.000 | 120 | 0.962 |
| Pro | P | CCC | GGG | 0 | 0.000 | 3 | 0.023 | 1 | 0.167 | 2 | 0.016 |

| | | | | | | | | | | | |
|---|---|---|---|---|---|---|---|---|---|---|---|
| Pro | P | CCA | TGG | 3 | 0.923 | 27 | 0.207 | 0 | 0.000 | 27 | 0.216 |
| Pro | P | CCG | CGG | 3 | 0.923 | 373 | 2.853 | 23 | 3.833 | 350 | 2.806 |
| Thr | T | ACT | AGT | 6 | 2.182 | 52 | 0.177 | 29 | 0.294 | 23 | 0.118 |
| Thr | T | ACC | GGT | 0 | 0.000 | 723 | 2.465 | 365 | 3.706 | 358 | 1.838 |
| Thr | T | ACA | TGT | 3 | 1.091 | 173 | 0.590 | 0 | 0.000 | 173 | 0.888 |
| Thr | T | ACG | CGT | 2 | 0.727 | 225 | 0.767 | 0 | 0.000 | 225 | 1.155 |
| Ala | A | GCT | AGC | 15 | 2.308 | 1126 | 2.519 | 1101 | 3.894 | 25 | 0.152 |
| Ala | A | GCC | GGC | 0 | 0.000 | 362 | 0.810 | 0 | 0.000 | 362 | 2.204 |
| Ala | A | GCA | TGC | 8 | 1.231 | 27 | 0.060 | 1 | 0.004 | 26 | 0.158 |
| Ala | A | GCG | CGC | 3 | 0.462 | 273 | 0.611 | 29 | 0.103 | 244 | 1.486 |
| Tyr | Y | TAT | ATA | 0 | 0.000 | 242 | 0.263 | 2 | 0.006 | 240 | 0.403 |
| Tyr | Y | TAC | GTA | 9 | 2.000 | 1601 | 1.737 | 650 | 1.994 | 951 | 1.597 |
| His | H | CAT | ATG | 0 | 0.000 | 467 | 0.848 | 17 | 0.343 | 450 | 0.897 |
| His | H | CAC | GTG | 7 | 2.000 | 635 | 1.152 | 82 | 1.657 | 553 | 1.103 |
| Gln | Q | CAA | TTG | 3 | 0.400 | 14 | 0.030 | 1 | 0.002 | 13 | 0.500 |
| Gln | Q | CAG | CTG | 12 | 1.600 | 913 | 1.970 | 874 | 1.998 | 39 | 1.500 |
| Asn | N | AAT | ATT | 0 | 0.000 | 501 | 1.876 | 497 | 1.949 | 4 | 0.333 |
| Asn | N | AAC | GTT | 7 | 2.000 | 33 | 0.124 | 13 | 0.051 | 20 | 1.667 |
| Lys | K | AAA | TTT | 6 | 0.750 | 259 | 0.442 | 196 | 0.363 | 63 | 1.385 |
| Lys | K | AAG | CTT | 10 | 1.250 | 913 | 1.558 | 885 | 1.637 | 28 | 0.615 |
| Asp | D | GAT | ATC | 0 | 0.000 | 611 | 0.599 | 536 | 0.837 | 75 | 0.198 |
| Asp | D | GAC | GTC | 7 | 2.000 | 1428 | 1.401 | 745 | 1.163 | 683 | 1.802 |
| Glu | E | GAA | TTC | 4 | 0.889 | 1352 | 0.725 | 624 | 0.460 | 728 | 1.430 |
| Glu | E | GAG | CTC | 5 | 1.111 | 2379 | 1.275 | 2089 | 1.540 | 290 | 0.570 |
| Cys | C | TGT | ACA | 0 | 0.000 | 394 | 1.003 | 394 | 1.003 | 0 | 0.000 |
| Cys | C | TGC | GCA | 28 | 2.000 | 392 | 0.997 | 392 | 0.997 | 0 | 0.000 |
| Trp | W | TGG | CCA | 5 | 1.000 | 1071 | 1.000 | 950 | 1.000 | 121 | 1.000 |
| Arg | R | CGT | ACG | 6 | 2.400 | 434 | 0.633 | 369 | 0.638 | 65 | 0.607 |

| | | | | | | | | | | | |
|---|---|---|---|---|---|---|---|---|---|---|---|
| Arg | R | CGC | GCG | 0 | 0.000 | 722 | 1.054 | 721 | 1.247 | 1 | 0.009 |
| Arg | R | CGA | TCG | 2 | 0.800 | 256 | 0.374 | 4 | 0.007 | 252 | 2.351 |
| Arg | R | CGG | CCG | 1 | 0.400 | 901 | 1.315 | 749 | 1.296 | 152 | 1.418 |
| Ser | S | AGT | ACT | 0 | 0.000 | 176 | 0.454 | 9 | 0.068 | 167 | 0.654 |
| Ser | S | AGC | GCT | 5 | 1.154 | 674 | 1.740 | 396 | 2.996 | 278 | 1.089 |
| Arg | R | AGA | TCT | 3 | 1.200 | 853 | 1.245 | 803 | 1.389 | 50 | 0.467 |
| Arg | R | AGG | CCT | 3 | 1.200 | 945 | 1.379 | 822 | 1.422 | 123 | 1.148 |
| Gly | G | GGT | ACC | 0 | 0.000 | 104 | 0.122 | 2 | 0.003 | 102 | 1.033 |
| Gly | G | GGC | GCC | 12 | 2.087 | 775 | 0.908 | 773 | 1.025 | 2 | 0.020 |
| Gly | G | GGA | TCC | 7 | 1.217 | 660 | 0.774 | 369 | 0.489 | 291 | 2.947 |
| Gly | G | GGG | CCC | 4 | 0.696 | 1874 | 2.196 | 1874 | 2.484 | 0 | 0.000 |

**Table S14** *Pearson correlation coefficients of correlations between the normalized relative frequencies of tRNA isotypes and RSCU among the 390 empirical MHC-I exon 3 sequences and the mean Pearson correlation coefficients between the normalized relative frequencies of tRNA isotypes and RSCU among the 1,000 data sets simulated under the assumption of no selection on synonymous variation (Mean sim.). C.I. lower and C.I. upper indicate the boundaries of 95% confidence intervals for the coefficients obtained with the simulated data sets, and the p-values indicate the proportion of coefficients with the simulated data sets more extreme than the value observed with the empirical data set. The N tRNA species column shows the number of codons matched by tRNA isotypes for each amino acid.*

| Amino acid | | N tRNA species | Pearson correlation coefficient | | | | |
|---|---|---|---|---|---|---|---|
| | | | Empirical | Mean sim. | C.I. lower | C.I. upper | P-value |
| Phe | F | 1 | NA | NA | NA | NA | NA |
| Leu | L | 5 | 0.43 | 0.28 | 0.21 | 0.35 | 0 |
| Ile | I | 3 | -0.31 | -0.11 | -0.20 | -0.02 | 0 |
| Met | M | 1 | NA | NA | NA | NA | NA |
| Val | V | 4 | 0.30 | 0.14 | 0.03 | 0.23 | 0 |
| Ser | S | 4 | 0.12 | -0.24 | -0.28 | -0.20 | 0 |
| Pro | P | 3 | -0.26 | -0.33 | -0.48 | -0.16 | 0.17 |
| Thr | T | 3 | -1.00 | -0.55 | -0.64 | -0.46 | 0 |
| Ala | A | 3 | 0.80 | -0.69 | -0.75 | -0.63 | 0 |
| Tyr | Y | 1 | NA | NA | NA | NA | NA |
| His | H | 1 | NA | NA | NA | NA | NA |
| Gln | Q | 2 | 1 | 1 | 1 | 1 | 0 |
| Asn | N | 1 | NA | NA | NA | NA | NA |
| Lys | K | 2 | 1 | 1 | 1 | 1 | 0 |
| Asp | D | 1 | NA | NA | NA | NA | NA |
| Glu | E | 2 | 1 | 1 | 1 | 1 | 0.016 |
| Cys | C | 1 | NA | NA | NA | NA | NA |
| Trp | W | 1 | NA | NA | NA | NA | NA |
| Arg | R | 5 | -0.32 | -0.38 | -0.43 | -0.32 | 0.03 |
| Gly | G | 3 | -0.73 | 0.10 | 0.01 | 0.20 | 0 |

**Table S15** *Pearson correlation coefficients of correlations between the normalized relative frequencies of tRNA isotypes and RSCU among the 390 MHC-I exon 3 sequences in the 57 sites where codon usage was under strong purifying selection and the 30 sites where purifying selection was relaxed, and the difference between them. The N tRNA species column shows the number of codons matched by tRNA isotypes for each amino acid.*

| Amino acid | | N tRNA species | Pearson correlation coefficient | | |
|---|---|---|---|---|---|
| | | | Purifying sel. | Relaxed sel. | Difference |
| Phe | F | 1 | NA | NA | NA |
| Leu | L | 5 | 0.32 | 0.52 | -0.20 |
| Ile | I | 3 | -0.30 | -0.37 | 0.07 |
| Met | M | 1 | NA | NA | NA |
| Val | V | 4 | 0.77 | -0.36 | 1.13 |
| Ser | S | 4 | -0.26 | 0.57 | -0.83 |
| Pro | P | 3 | -0.50 | -0.24 | -0.26 |
| Thr | T | 3 | 0.97 | -1.00 | 1.97 |
| Ala | A | 3 | 0.90 | -0.82 | 1.72 |
| Tyr | Y | 1 | NA | NA | NA |
| His | H | 1 | NA | NA | NA |
| Gln | Q | 2 | 1 | 1 | 0 |
| Asn | N | 1 | NA | NA | NA |
| Lys | K | 2 | 1 | -1 | 2 |
| Asp | D | 1 | NA | NA | NA |
| Glu | E | 2 | 1 | -1 | 2 |
| Cys | C | 1 | NA | NA | NA |
| Trp | W | 1 | NA | NA | NA |
| Arg | R | 5 | -0.15 | -0.60 | 0.45 |
| Gly | G | 3 | -0.60 | -0.14 | -0.46 |

**Table S16** *Generalized linear models of the effect of synonymous nucleotide variation on life span. Synonymous nucleotide variation was quantified as the mean number of synonymous nucleotide changes in pairwise comparisons among the sequences in each individual. The table shows the results of the final model for both sexes, which included sex as a fixed factor and the interaction 'mean number of synonymous nucleotide changes × sex', and of separate analyses for females and males.*

| **Life span** | | | | |
|---|---|---|---|---|
| **Both sexes** | **Estimate** | **Std. error** | **t-value** | **Pr(>\|t\|)** |
| Intercept | 1.62 | 0.56 | 2.90 | 0.0043 |
| Mean no. syn. changes | -0.058 | 0.084 | -0.70 | 0.49 |
| Sex (male) | -1.61 | 0.91 | -1.77 | 0.078 |
| Mean no. syn. changes × sex (male) | 0.24 | 0.13 | 1.80 | 0.074 |
| **Females** | | | | |
| Intercept | 1.62 | 0.64 | 2.54 | 0.013 |
| Mean no. syn. changes | -0.058 | 0.095 | -0.61 | 0.54 |
| **Males** | | | | |
| Intercept | 0.0054 | 0.57 | 0.01 | 0.99 |
| Mean no. syn. changes | 0.18 | 0.083 | 2.21 | 0.030 |

**Table S17** *Generalized linear models of the effect of synonymous nucleotide variation on lifetime number of fledged offspring. Synonymous nucleotide variation was quantified as the mean number of synonymous nucleotide changes in pairwise comparisons among the sequences in each individual. The table shows the results of the final model for both sexes, which included sex as a fixed factor and the interaction 'mean number of synonymous nucleotide changes × sex', and of separate analyses for females and males.*

| **Lifetime number of fledged offspring** | | | | |
|---|---|---|---|---|
| **Both sexes** | **Estimate** | **Std. error** | **t-value** | **Pr(>\|t\|)** |
| Intercept | 2.45 | 0.79 | 3.09 | 0.0023 |
| Mean no. syn. changes | -0.015 | 0.12 | -0.12 | 0.90 |
| Sex (male) | -2.25 | 1.28 | -1.76 | 0.080 |
| Mean no. syn. changes × sex (male) | 0.38 | 0.19 | 1.99 | 0.048 |
| **Females** | | | | |
| Intercept | 2.45 | 0.76 | 3.23 | 0.0017 |
| Mean no. syn. changes | -0.015 | 0.11 | -0.13 | 0.90 |
| **Males** | | | | |
| Intercept | 0.20 | 1.05 | 0.19 | 0.85 |
| Mean no. syn. changes | 0.36 | 0.16 | 2.33 | 0.022 |

**Table S18** *Generalized linear model of the effect of synonymous nucleotide variation on lifetime number of fledged offspring adjusted for life span. Synonymous nucleotide variation was quantified as the mean number of synonymous nucleotide changes in pairwise comparisons among the sequences in each individual. The table shows the results of the final model for both sexes, which included sex as a fixed factor.*

| **Lifetime number of fledged offspring (adj. for life span)** | | | | |
|---|---|---|---|---|
| **Both sexes** | **Estimate** | **Std. error** | **t-value** | **Pr(>\|t\|)** |
| Intercept | 0.48 | 0.40 | 1.21 | 0.23 |
| Life span | 0.30 | 0.020 | 15.13 | 3.07E-33 |
| Mean no. syn. changes | 0.10 | 0.059 | 1.77 | 0.079 |
| Sex (male) | 0.32 | 0.076 | 4.24 | 3.66E-05 |

**Table S19** *Generalized linear model of the effect of number of different MHC-I sequences and mean amino acid p-distance on life span. Mean amino acid p-distance was quantified as the mean proportion of amino acid changes in pairwise comparisons among the sequences in each individual. The table shows the results of the full model, which included sex as a fixed factor and the interactions 'number of different MHC-I sequences × sex' and 'mean amino acid p-distance × sex'.*

| **Life span** | | | | |
|---|---|---|---|---|
| **Both sexes** | **Estimate** | **Std. error** | **t-value** | **Pr(>\|t\|)** |
| Intercept | 1.47 | 0.54 | 2.70 | 0.0077 |
| No. different MHC-I sequences | 0.0043 | 0.019 | 0.23 | 0.82 |
| Mean a.a. p-distance | -2.06 | 3.63 | -0.57 | 0.57 |
| Sex (male) | 0.30 | 0.94 | 0.32 | 0.75 |
| No. different MHC-I sequences × sex (male) | -0.0089 | 0.027 | -0.33 | 0.74 |
| Mean a.a. p-distance × sex (male) | -1.20 | 6.33 | -0.19 | 0.85 |

**Table S20** *Generalized linear model of the effect of number of different MHC-I sequences and mean amino acid p-distance on lifetime number of fledged offspring. Mean amino acid p-distance was quantified as the mean proportion of amino acid changes in pairwise comparisons among the sequences in each individual. The table shows the results of the full model, which included sex as a fixed factor and the interactions 'number of different MHC-I sequences × sex' and 'mean amino acid p-distance × sex'.*

| **Lifetime number of fledged offspring** | | | | |
|---|---|---|---|---|
| **Both sexes** | **Estimate** | **Std. error** | **t-value** | **Pr(>\|t\|)** |
| Intercept | 2.47 | 0.77 | 3.22 | 0.0016 |
| No. different MHC-I sequences | 0.018 | 0.026 | 0.69 | 0.49 |
| Mean a.a. p-distance | -2.54 | 5.11 | -0.50 | 0.62 |
| Sex (male) | 1.72 | 1.31 | 1.31 | 0.19 |
| No. different MHC-I sequences × sex (male) | -0.036 | 0.038 | -0.95 | 0.35 |
| Mean a.a. p-distance × sex (male) | -6.62 | 8.87 | -0.75 | 0.46 |

**Table S21** *Generalized linear model of the effect of number of different MHC-I sequences and mean amino acid p-distance on lifetime number of fledged offspring adjusted for life span. Mean amino acid p-distance was quantified as the mean proportion of amino acid changes in pairwise comparisons among the sequences in each individual. The table shows the results of the full model, which included sex as a fixed factor and the interactions 'number of different MHC-I sequences × sex' and 'mean amino acid p-distance × sex'.*

| **Lifetime number of fledged offspring (adj. for life span)** | | | | |
|---|---|---|---|---|
| **Both sexes** | **Estimate** | **Std. error** | **t-value** | **Pr(>\|t\|)** |
| Intercept | 0.76 | 0.52 | 1.48 | 0.14 |
| Life span | 0.30 | 0.020 | 14.74 | 6.38E-32 |
| No. different MHC-I sequences | 0.010 | 0.017 | 0.57 | 0.57 |
| Mean a.a. p-distance | 1.93 | 3.34 | 0.58 | 0.56 |
| Sex (male) | 1.12 | 0.86 | 1.31 | 0.19 |
| No. different MHC-I sequences × sex (male) | -0.020 | 0.025 | -0.80 | 0.42 |
| Mean a.a. p-distance × sex (male) | -3.67 | 5.76 | -0.64 | 0.52 |

**Table S22** *Log likelihood, number of parameters, and AIC values for 12 substitution models in PhyML. The models are listed by AIC values. The basic models were JC69, HKY85, and generalized time-reversible model (GTR), each with additional estimation of gamma shape parameters (G), proportion of invariable sites (I), or both (G + I). The number of parameters in each model was derived as: no. parameters in model = (2 × no. sequences – 3) + no. parameters in substitution model + extra parameters, where no. sequences = 390 and no. substitution model parameters are: JC = 0 parameters, HKY = 4 parameters, GTR = 8 parameters. Extra parameters are: Invariable sites = +1 parameter, and gamma rates = +1 parameter. AIC values were calculated for each model by the formula: AIC = 2k – 2 × log(L), where log(L) is the log-likelihood ratio value obtained for each model.*

| Substitution model | log(L) | Parameters | AIC |
|---|---|---|---|
| GTR | -7059.73 | 787 | 15693.47 |
| GTR + I | -7060.23 | 788 | 15696.46 |
| GTR + G + I | -7093.55 | 789 | 15765.10 |
| HKY + I | -7105.83 | 784 | 15779.65 |
| HKY | -7125.53 | 783 | 15817.07 |
| GTR + G | -7121.32 | 788 | 15818.65 |
| HKY + G | -7147.98 | 784 | 15863.95 |
| HKY + G + I | -7162.26 | 785 | 15894.52 |
| JC + I | -7293.09 | 780 | 16146.18 |
| JC + G + I | -7307.02 | 781 | 16176.03 |
| JC | -7315.96 | 779 | 16189.92 |
| JC + G | -7329.97 | 780 | 16219.94 |

**Table S23** *3x4 table of base frequencies by (within codon) nucleotide position among the 390 Great Reed Warbler MHC-I exon 3 sequences output by the M2a and M8 models in codeml.*

| | T | C | A | G |
|---|---|---|---|---|
| Position 1 | 0.20607 | 0.21438 | 0.20227 | 0.37728 |
| Position 2 | 0.21789 | 0.14612 | 0.33445 | 0.30153 |
| Position 3 | 0.15912 | 0.36004 | 0.10813 | 0.37271 |

**Table S24** *Relative probabilities of synonymous codon variants calculated as products of the base frequencies by nucleotide position in Table S23. The values in this table were used as weights in the simulation of synonymous DNA sequences reflecting the null hypothesis that variation in codon usage is due to mutation bias alone.*

| Amino acid | | Codon | Freq 3x4 | Amino acid | | Codon | Freq 3x4 |
|---|---|---|---|---|---|---|---|
| Phe | F | TTT | 0.00714458 | Tyr | Y | TAT | 0.01096657 |
| | | TTC | 0.01616601 | | | TAC | 0.02481400 |
| Leu | L | TTA | 0.00485510 | STOP | X | TAA | - |
| | | TTG | 0.01673490 | | | TAG | - |
| Leu | L | CTT | 0.00743270 | His | H | CAT | 0.01140881 |
| | | CTC | 0.01681792 | | | CAC | 0.02581465 |
| | | CTA | 0.00505089 | Gln | Q | CAA | 0.00775286 |
| | | CTG | 0.01740975 | | | CAG | 0.02672308 |
| Ile | I | ATT | 0.00701283 | Asn | N | AAT | 0.01076434 |
| | | ATC | 0.01586790 | | | AAC | 0.02435642 |
| | | ATA | 0.00476557 | Lys | K | AAA | 0.00731491 |
| Met | M | ATG | 0.01642630 | | | AAG | 0.02521353 |
| Val | V | GTT | 0.01308055 | Asp | D | GAT | 0.02007797 |
| | | GTC | 0.02959728 | | | GAC | 0.04543031 |
| | | GTA | 0.00888888 | Glu | E | GAA | 0.01364398 |
| | | GTG | 0.03063883 | | | GAG | 0.04702903 |
| Ser | S | TCT | 0.00479125 | Cys | C | TGT | 0.00988713 |
| | | TCC | 0.01084115 | | | TGC | 0.02237155 |
| | | TCA | 0.00325590 | STOP | X | TGA | - |
| | | TCG | 0.01122265 | Trp | W | TGG | 0.02315882 |
| Pro | P | CCT | 0.00498447 | Arg | R | CGT | 0.01028584 |
| | | CCC | 0.01127833 | | | CGC | 0.02327371 |
| | | CCA | 0.00338719 | | | CGA | 0.00698974 |
| | | CCG | 0.01167522 | | | CGG | 0.02409272 |
| Thr | T | ACT | 0.00470290 | Ser | S | AGT | 0.00970480 |
| | | ACC | 0.01064123 | | | AGC | 0.02195901 |
| | | ACA | 0.00319586 | Arg | R | AGA | 0.00659490 |
| | | ACG | 0.01101570 | | | AGG | 0.02273176 |
| Ala | A | GCT | 0.00877199 | Gly | G | GGT | 0.01810169 |
| | | GCC | 0.01984834 | | | GGC | 0.04095860 |
| | | GCA | 0.00596101 | | | GGA | 0.01230100 |
| | | GCG | 0.02054681 | | | GGG | 0.04239995 |

**Fig. S1** *Ribbon diagram of the protein structure predicted by AlphaFold 3 from the consensus amino acid sequence derived from the 390 Great Reed Warbler MHC-I exon 3 sequences. Numbers indicate the amino acid sites in each β-sheet and α-helix domain (Table S7). Image generated at alphafoldserver.com.*

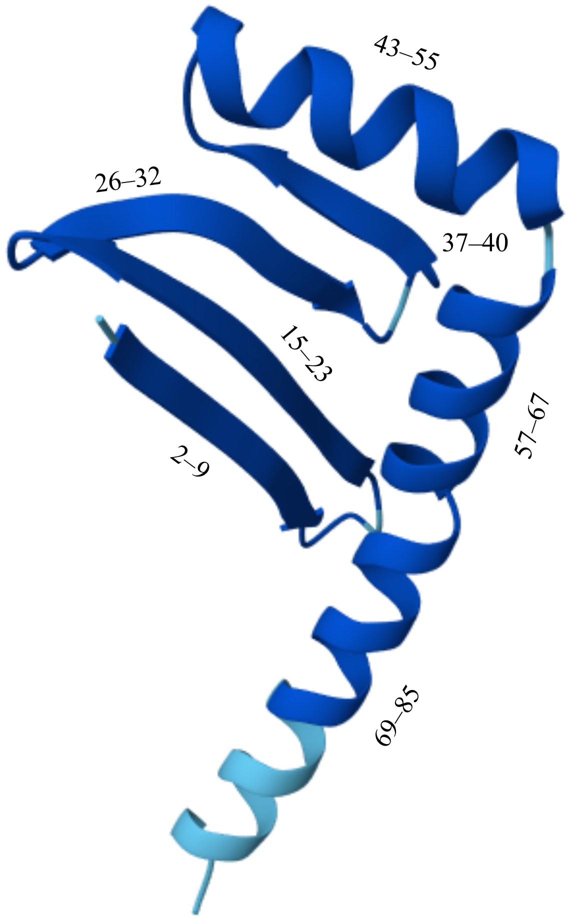

**Fig. S2** *Distribution of the mean number of synonymous nucleotide changes in pairwise comparisons among the sequences in each individual sample.*

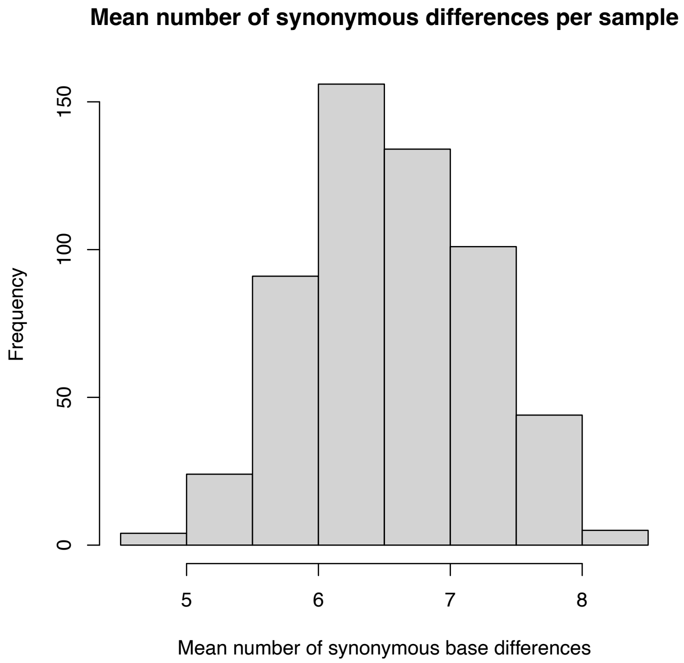

**Fig. S3** *The relationship between synonymous nucleotide variation and offspring fledging success (lifetime number of fledged offspring adjusted for life span) in adult Great Reed Warblers. In this plot, offspring fledging success was calculated as residuals from a linear regression between lifetime number of fledged offspring and life span. These residuals were approximately normally distributed, and model predictions (indicated by the lines) were obtained from a linear regression model of the effects of mean number of synonymous nucleotide changes on these residuals including sex as fixed factor. Open circles and dashed line indicate females. Black triangles and solid line indicate males.*

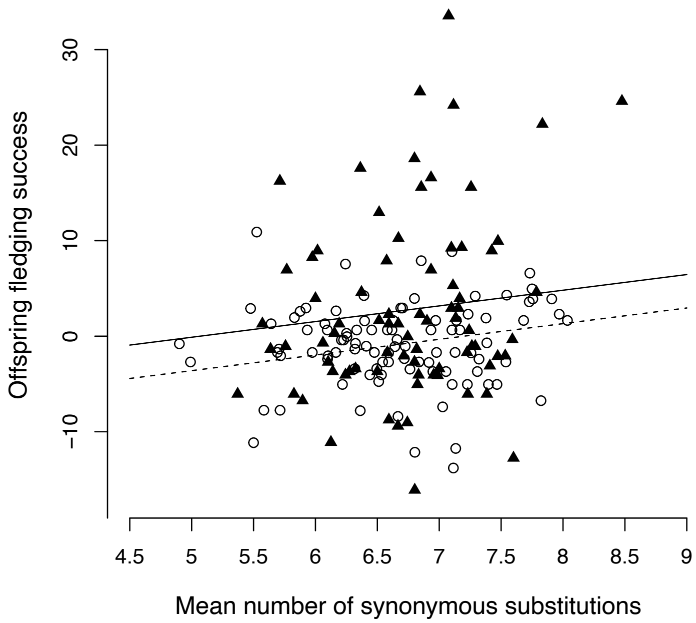
